# Identification of Prospective PETases Across Prokaryotes Using an *in silico* Approach

**DOI:** 10.1101/2025.08.14.670228

**Authors:** Yashkumar Rathod, Sumit Biswas

## Abstract

**Background:** Plastics count as one of the most potent threats to the habitats and survival of global flora and fauna. Reports keep accumulating globally about the ever-exploding load of plastic wastes, but the need and economics of multiple industrial and household processes compels the production of more plastic materials. It has always been imperative to look for natural sources of degradation of plastic. The identification of plastic-degrading microbes, therefore remains a major focus of the microbial fraternity. While the discoveries of *Ideonella sakaiensis* or later, *Rhizobacter gummiphilus* were more out of providence, the structure determination of the enzyme responsible for PET degradation does provide a fillip to efforts towards identification of more such prokaryotic entities.

**Results:** In this work, a comprehensive profiling of prokaryotic sequences has been undertaken to look for the presence of similar plastic-degradation abilities across the kingdom. The identification of twenty-seven such ‘hits’ across different bacterial species led us to believe in the natural diversity of plastic-degradation enzymes. Moreover, there seems to be conservation of the structural motif that renders such ability as has been observed from the constructed models and analysis of their interfaces. Docking of BHET, one of the key products of PET, against these 27 entities showed considerable interactions with the above and pointed towards the possible roles of these bacteria as natural plastic degradation models. Eight of these proteins have very close similarity in binding interactions and surface properties to the PETase from *I. sakaiensis* and were shortlisted as prospective candidates.

**Conclusions:** Of these eight, three PETases from *Halopseudomonas pertucinogena Halopseudomonas bauzanensis* and *Ketobacter sp.* revealed significant similarity in structure and conformational stability to the PETase from *I. sakaiensis* as was evident from the analysis of their molecular dynamics parameters. Principal Component Analysis and the free energy landscape during binding to BHET also validated the hypothesis and these three PETases could be immediately explored for possible plastic degradation activity.

## Introduction

The world is burdened by a burgeoning load of plastics and the accumulation of these contaminants in the ecosystem is a potent threat to life and livelihood [1, 2]. The array of these plastic pollutants, of different sizes and types is also mind-boggling. While the load of plastic pollution keeps increasing, driven by global demand and eventual production, means and methods of plastic recycling are few and far in between and ineffective on a broader perspective. The staggering range of the burden is often disturbing. The yearly production of plastic waste has been projected to reach 3.4 billion tonnes by 2050, according to the study done by World Bank [3].

In an effort to limit the load of plastic pollutants, degradation of plastics using microbial sources has been the subject of study over the past few decades. The polymerization of plastic is effected by means of weak forces like van der Waals interactions, electrostatic forces as well as hydrogen bonds. etc. Microbes are known to degrade plastics in a sequential manner. Broadly, this can be simplified as physical or chemical deterioration, followed by breakdown of polymers which are then shipped to the cytoplasm of the microbes for metabolic breakdown [3].

One of the biggest sources of the pollution and perhaps the most commonly used of plastics is Polyethylene Terephthalate (PET). Microbial biodegradation of PET has been mooted for long and a few breakthroughs have been achieved over the years. Enzymes from *Thermobifida fusca* were reported from the early years of this century to be effective in the degradation of PET[4]. Lipases from *Candida antarctica* [4] and cutinases from *Humicola insolens* [5] have also been known to display PET degradation, and some of these have been shown to have synergistic effects in elevating the efficacy and reducing the time of the degradation process [6].

*Ideonella sakaiensis*, a plastic-degrading bacterium, was first discovered in 2016 [7]. *I. sakaiensis* is a Gram negative, aerobic, rod-shaped bacterium belonging to the genus Ideonella and family Comamonadaceae [8]. Originally isolated from a sediment sample taken outside of a plastic bottle recycling facility in Sakai, the bacterium also uses PET as a sole carbon and energy source for growth. The PETase enzyme found in the bacteria aids in the degradation of the ester bond of PET. IsPETase (PETase from *I. sakaiensis*) has demonstrated the highest plastic degradation activity compared to all other bacterial enzymes despite the fact that similar enzymes have been discovered in a number of other organisms.

The mechanism of PET degradation by the IsPETase enzyme [9] has been studied and reported by numerous groups. The PETase enzymes showed the presence of a conserved, PET-binding active site. Ser (160), Asp (206) and His (237) form a highly conserved catalytic triad that aids in the stable binding with PET. Several mutations were introduced into the PETase enzyme of *Ideonella sakaiensis*, and wild-type bacteria were used for comparison. [6] has also reported that mutant IsPETase R280A is 22.4% more effective than the wild type. According to the report, these variants were more effective at higher temperatures and their plastic degradation rate was significantly higher compared to the wild type, but they are not effectively utilised for refuse degradation [10]. However, wild-type bacteria have certain limitations that prevent large-scale application. IsPETase in its natural state is exceedingly unstable and therefore, IsPETase have also been modified to degrade PET completely at higher temperatures [11].

The mechanism and high-resolution crystal structure of IsPETase [12] showed that the enzyme has a hydrophobic cavity that binds to PET polymers and breaks the PET into monomers like bis(2-hydroxyethyl)-TPA (BHET) and mono-(2-hydroxyethyl) terephthalic acid (MHET) 13. IsPETase though is incapable of hydrolyzing MHET to terephthalic acid (TPA) and ethylene glycol (EG). Very recently, [14] reported that PETase from *Rhizobacter gummiphilus* has comparable binding affinity for PET and hydrolysis activity at moderate temperatures. With very few substitutions in the conserved domain, RgPETase was found to be remarkably similar to IsPETase. These are though, the only two instances of discovery of PET degrading bacteria till date.

The discovery of plastic degradation abilities in prokaryotic species is a slow process, and discoveries are few and far in between. With the knowledge of the available enzymes and the couple of structures that have been solved, we therefore strived to predict the presence of such plastic degradation abilities in other bacterial species. The search was manifold, both at the level of the sequences and the conservation of the degradation domain. The hits were subjected to *in silico* verification using structural parameters and docking tools. In this investigation, BHET was chosen as the substrate both due to its structural similarity to the nucleus of PET and it’s ability to hydrolyze into TPA and EG monomers. The predicted structures and the enzymes shed light on the variety of plastic degradation domains from across the prokaryotes, and also demonstrate the range of alternatives that could be explored in addition to *I. sakaiensis* for plastic degradation mediated by bacteria.

## Methods

### 1. Sequence alignment and conserved domain identification

The enzyme polyethylene terephthalate hydrolase (PETase) of *I. sakaiensis* (PDB ID: 6ILW) [9] with resolution 1.57 Å was used as a target in this study. This structure was chosen on the basis of maximum completeness among the other structures of PETase available in the PDB. Sequences similar to those of IsPETase, from across all prokaryotes, were searched for using the Protein Blast tool of NCBI [15], with a selected threshold of 95% query coverage. 27 sequences were thus obtained and aligned against the sequence of 6ILW (supplementary table 1 and figure S6). The obtained sequences were subjected to domain identification using MotifScan [16], using motif sources from Pfam [17] and HAMAP [18]. Based on the lowest e-value, domains were identified on the basis of their conserved nature.

Sequences of the domains (supplementary figure S8) thus retrieved, were aligned among the 27 hits and the total number of mutations (substitutions and indels) were identified. IsPETase structure was submitted to the Consurf web server [19]. The web server identified conserved amino acids based on the sequence alignment and the conservation score was fixed in the range of 1 to 9, with various colours designated to each value, and the protein structure was represented by colour patches based on their conservation values (supplementary figure S4).

### 2. Protein Interactome

To understand the nature of the interactions of the PETase protein with other proteins in the cell, the STRING database (https://string-db.org/) was used to create the protein-protein interactions (PPIs) network*. I. sakaiensis* was chosen as the query organism in the STRING database’s “search protein by name” function in order to navigate the database. The minimum required interaction score was set to “low confidence” so that an exhaustive search of the proteins in the organism is performed. The size of the interactions was kept at “not >50 interactions” in the first and second shell to have a fair idea of the interacting proteins with PETase.

### 3. Homology Modelling

The structure of the 27 entries listed before were searched for in the PDB, and revealed that only one of the entries (7DZT, *Rhizobacter sp.)* had a deposited structure. To compensate for the lack of crystal structure, the rest of the 26 entries were subjected to homology modelling using Maestro module [20] of Schrödinger release 2023.1 with customized settings for structural prediction. Option of “One Target Multiple Templates” was selected, followed by selection of top ten similar structures for the models. PROCHECK [21]and ERRAT [22]were used to verify the quality of the predicted structures. PyMOL was used to view and depict the final structures. The modelled structures were aligned with an aim to understand deviations from the template structure of 6ILW.

### 4. Docking and molecular dynamics simulation studies

The ligand for docking at the PETase site was chosen to be the 3D conformer of bis (2-hydroxyethyl) terephthalate (BHET) retrieved from PubChem. The 3D structure-data files (SDF) format was utilised for docking analysis. The docked structures were analyzed for the lowest energy conformer, Root mean square deviation (RMSD) and hydrogen bond (HB) interaction.

#### Receptor Preparation

The basic receptor molecule was chosen to be the enzyme PET hydrolase of *I. sakaiensis* (PDB ID: 6ILW) with resolution 1.57 Å. The protein had a molecular weight of 28.6 kDa and contained 265 amino acids. The target was prepared by removing water, repairing broken side chains, adding polar hydrogens, and adding Gasteiger charges to balance surface charges. The same was done for all the other modelled structures obtained from homology modelling.

#### Binding Site Prediction

The active site of the enzyme was identified from literature and grid of dimensions 17.1648 x 14.3731 x 17.5487 Å was built around it, centered at 22.4021, 7.7184 and 10.6203 Å. A grid box was thereby constructed around subsite 1 (Tyr87, Met161, Trp185, Ile208) and subsite 2 (Thr88, Ala89, Trp159, Ile232, Asn233, Ser236, Ser238, Asn241, Asn244, Ser246 and Arg280), reported to be the active site of the enzyme. From the structure of IsPETase, it was reported that subsites 1 and 2 aid in the stable binding of ligand and in case of any substitution in the catalytic triad (Ser160, Asp206, His 237) the ligand’s binding affinity is altered [10]. A conserved serine hydrolase motif Gly-X1-Ser-X2-Gly (supplementary figure S8) is also a part of this catalytic triad, and the grid box was aimed at these regions for all the protein structures. Ligand docking of the BHET compound onto the enzyme targets was accomplished with the help of Autodock Vina in version 0.8 of the PyRx programme [23, 24]. Prepared molecule was loaded into the PyRx programme and grid box was set to the position center x: 22.4021, center y: 7.7184, center z: 10.6203, and size x: 17.1648, size y: 14.3731, size z: 17.5487.

#### Molecular Dynamics

Molecular Dynamics (MD) simulation was performed utilizing Gromacs 2021.3 through the command line interface [25]. Proteins were vacuum energy minimized before the actual MD procedure to obtain the optimized PDB structure. The most optimal PETase was identified from docking scores and RMSD, followed by subsequent 100 nanosecond molecular dynamics (MD) simulation, repeated over three iterations each. The simulation employed the Charmm36 forcefield. The protein topology file was constructed using the Gromacs software, whereas the ligand topology file was prepared using the CGenFF server [26]. For solvating the complex, the TIP3P water model was used in a 1 nm-sized dodecahedron box. To counterbalance the overall charge of the protein-ligand complex, an adequate amount of sodium (Na+) or chloride (Cl-) ions were added. The system’s energy minimisation was performed using the steepest descent algorithm for 50,000 iterations to resolve steric clashes.

The C-rescale and V-rescale coupling approach was used for equilibration in NVT and NPT ensemble simulations respectively, with positional restrains imposed on the protein and ligand to preserve its structural integrity during the equilibration phase. The structural coordinates of the system were saved for every 10 ps by GROMACS. Generated MD trajectories were analysed in order to calculate root mean square deviation (RMSD), root mean square fluctuation (RMSF) and to conduct Principal Component Analysis (PCA). The conformational stability, fluctuating residues, and compactness of the protein and the protein-BHET complexes were validated using the gmx rms, gmx rmsf, gmx gyrate modules of GROMACS, respectively. The Molecular Mechanics/Poisson-Boltzmann Surface Area (MMPBSA) approach was utilised to determine the relative binding free energy of protein-ligand complexes. The gmx_MMPBSA tool was employed to compute the binding free energies of protein-ligand complexes, using 1000 frames collected from the final 10 ns (90–100 ns) of the molecular dynamics simulation.

#### Principal Component Analysis (PCA)

Principal Component Analysis (PCA) was conducted in this study to examine the motion and compactness of the molecule during the simulation. The covariance matrix was constructed using the gromacs “gmx covar” module based on the previously determined trajectories. The eigen vectors and eigen values obtained from the covariance matrix were analysed using the “gmx anaeig” package. The eigen vector indicates the direction of motion, whereas the eigenvalues indicate the degree of motion along each major component. The data was stored in .xvg file, and a software application based on Python programming language was constructed for the purpose of data visualization.

## Results and Discussion

### Sequence Similarities and Domain Identification

The search for sequences similar to the IsPETase across prokaryotes yielded a total number of 169 hits. Some of these (11) were mutants of IsPETase which had been generated for functional characterization, and were excluded from the search results. Similarly, 56 mutants of PETase from *I. sakaiensis* were filtered to remove any redundancies. Of the remaining hits, the best matches were selected – a process which yielded 27 sequences across various prokaryotic genomes, having similarity to IsPETase. All the 27 sequences which were selected showed the presence of the conserved serine hydrolase motif (GWSXG), which has been reported to be necessary for the binding of MHET to the active site of IsPETase. ConSurfDB displayed multiple highly conserved areas which were highlighted using a colour-coded approach, with the hydrolase domain classified among the highly conserved sections (supplementary figures S4 and S5). The presence of the cysteines nearabout the active site for secondary disulphide bond formation, which has also been implicated for PETase activity, is evident among all these 27 sequences.

The 27 sequences thus obtained were checked for their sources and revealed the presence of these sequences across *Rhizobacter sp., Acidovorax sp., Ketobacter sp., Piscinibacter sp.*, etc. The source species for the 27 have been listed in supplementary table 1. The obtained sequences were aligned using Clustal Omega and displayed conservation as high as 97.72% in PETase (WP_085749752.1) from *R. gummiphilus* with 99% sequence coverage. This was in complete agreement with the work of Sagong et al., 2021 [28]. The lowest similarity which was considered was the sequence from *Halopseudomonas pertucinogena* with a sequence identity of 51.32% and 98% sequence coverage. Apart from the conserved serine hydrolase motif and the cysteine, the presence of a conserved methionine near the serine hydrolase motif was also a hallmark of the 27 sequences (supplementary figure S6).

Further, the sequence alignment revealed that all the sequences belonged to the type II category of PETases, except *Proteobacteria bacterium*. Type II PETases are typically characterized by additional disulphide linkages due to the conserved cysteine residues at positions 203 and 239. Type II PETases are more efficient in their plastic degradation abilities compared to type I PETases due to the additional disulphide bonds and certain substitutions in the subsite 2 (supplementary figure S6) [6].

Domain identification using MotifScan with the Pfam dataset revealed the presence of the esterase domain acetyl xylene esterase (AXE) across all the 27 sequences. The AXE domain was identified at position 123 to 155 in *I. sakaiensis* exhibiting an E value of 0.00049 and belonging to the pfam_FS family. The Pfam FS is a database of complete sequences, encompassing the full diversity of proteins, to reveal maximum identification. The AXE domain, identified by Pfam FS, is typified by a Ser at position 160, a Met at 161 and four consecutive Gly residues after the M161 (supplementary figure S7 and S8) and this was used for all further analyses.

In the analysis of 27 hits, the amino acid composition inside the domain exhibited alterations at 31% (283 out of 918) of the assessed sites, with substitutions being the primary alteration. Deletions were noted (supplementary table 2) in merely 1.5% (14 out of 918), whereas additions were negligible (2 out of 918).

### The Protein Interactome

The PPI network as created by the STRING database revealed a handful of proteins that are part of the IsPETase network. Some of these proteins have been implicated in critical processes in the associated pathway of plastic degradation. Remarkable among these was chlorogenate esterase (supplementary figure S3), which has synergistic effects with PETase in breaking down PET by catalysing the hydrolysis of MHET (mono(2-hydroxyethyl) terephthalate) into its two ecologically safe monomers, terephthalate and ethylene glycol. Chlorogenate esterase (MHETase) interacts with both subsites 1 and 2 for the subsequent degradation of MHET [7, 30]. In the absence of MHETase, PET is transformed into MHET, TPA, and EG but no further degradation of MHET happens. TPA and EG serve as more readily useable carbon sources, but MHET necessitates additional degradation to be transformed into TPA and EG. The other proteins involved in the network are basic functional proteins like topoisomerases or serine kinases with one membrane protein also finding its place.

### Similarities in Structures of Predicted PETases

Given the recent discovery of PETase, not many structures were available and it was difficult to determine the structures from defined templates. To confirm the similarities in the sequence and the domains, the hits were modelled against the template of the complete structure of IsPETase. The structure of RgPETase from *Rhizobacter gummiphilus.* was available from the PDB with the id. 7DZT. The homology models for the rest of the 26 hits were obtained using the Maestro module of Schrödinger and validated using ERRAT and Verify3D. Assessing the stereochemical properties using PROCHECK, the Ramachandran Plot for the modelled structures (from PROCHECK) also verified (supplementary table 3) the structural integrities (more than 94% in allowed regions for all entries except two where 90% were in allowed regions).

The structures were visualized in PyMOL and superimposed with the template structure (6ILW) to reveal striking similarities in their overall architecture, as shown in figure 1. The similarities in the predicted structures were also checked using GraSR [29] to verify the visual output from PyMOL. GraSR constructs a graph based on inter-residue distances derived from the tertiary structure. The output is generated based on the predictions from a deep graph neural network. The obtained T_M_ value represents the similarity of the structure on a scale of 0 to 1, with 1 being the most similar. For our predicted structures, the T_M_ for all the models were above 0.9, with the closest being the structure from *Acidovorax delafieldii*. This was indicative of strong structural similarity of the predicted structures with that of *Is*PETase. The T_M_ values have been listed in supplementary table 4.

**Figure 1:**
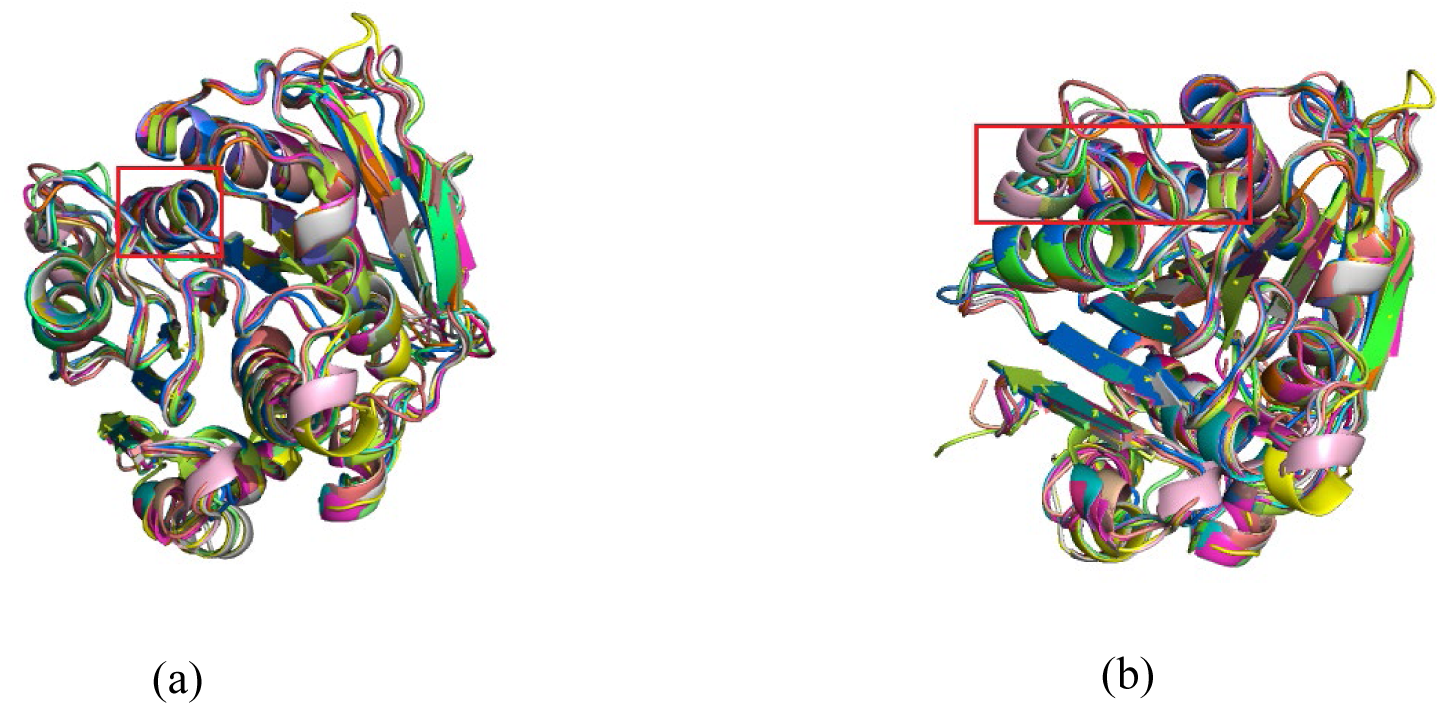
Superimposition of modelled structures with IsPETase revealed similarity in architecture across the AXE domains (highlighted in red box) and also in the overall structure (a) Anterior face, (b) Posterior face

### Heteromolecular associations of PETase-BHET complex

The top docking clusters of PETase bound to BHET showed engagement of BHET in the structure near the binding pocket encompassing the serine hydrolase motif across the 27 queries. The presence of a methionine at the docking site was notable in terms of its possible role in the binding of BHET, very much in accordance with previous report [30].

The most favorable binding pose of PETase-BHET complex showed stable binding with a docking score of −5.5 kcal/mol. The chemical structure of BHET and its heteromolecular associations with PETase have been demonstrated in figure 2. Detailed analysis of the binding pose indicated a strong association of PETase with BHET, stabilized by several hydrophobic interactions and hydrogen bonds. S 160, D 206, and H 237 were present at the catalytic site for IsPETase, and the same site was discovered to be highly conserved across other PETases.

**Figure 2:**
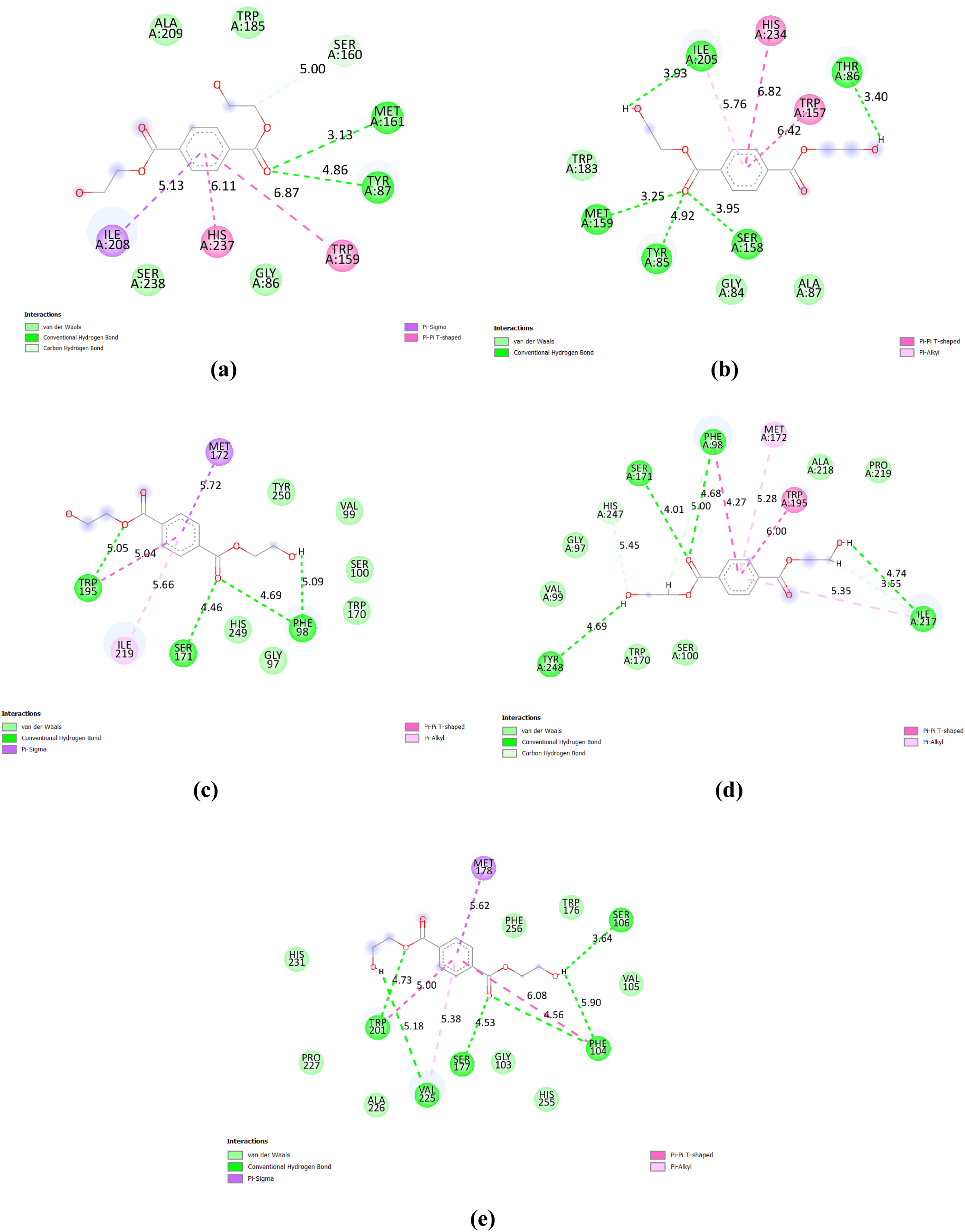
The protein-ligand interaction picture depicts the binding interactions between the ligand and particular amino acid residues in the protein’s active region. Essential interactions, including hydrogen bonds, hydrophobic contacts, and π-π stacking, are illustrated, emphasising the ligand’s binding mechanism and the stability of the complex. a) PETase from *Ideonella sakaiensis* b) PETase from *Rhizobacter gummiphilus*. c) PETase from *Halopseudomonas pertucinogena* d) PETase from *Halopseudomonas bauzanensis* e) PETase from *Ketobacter sp*.

**Table 1:** Hydrogen bonds and Hydrophilic interactions.

| Bacteria Name | Hydrogen bonds | Hydrophobic interactions |
| --- | --- | --- |
| *Ideonella sakaiensis* | Tyr (87), Met (161), Ile (208) | Trp (159), Ile (208), His (237) |
| *Rhizobacter gummiphilus* | Tyr (85), Thr (86), Ser (158), Met (159), Ile (205) | Trp (157), Ile (205), His (234) |
| *Marinobacter daqiaonensis* | Ala (106), Ser (107), Ser (178), Tyr (257) | Phe (105), Ile (226) |
| *Halopseudomonas pertucinogena* | Phe (98), Ser (171), Trp (195) | Ile (219), Trp(195) |
| *Pseudomonas zhaodongensis* | Tyr (93), Ser (95), Ser (166) | Met (167), Tyr (190), Val (214) |
| *Halopseudomonas bauzanensis* | Phe(98), Ser (171), Ile(217), Tyr (248) | Phe (98), Ile (217) |
| *Ketobacter sp.* | Phe (104), Ser (106), Ser (177), Trp(201), Val(225) | Phe(104), Trp (201), Val (225) |
| *Burkholderia sp.* | Gly (106) | Ala (109) |
| *Aquabacterium sp.* | Tyr (86), His (158), Trp (184), Ile (207) | Met (160) |
| *Candidatus muproteobacteria bacterium* | Tyr (77), Ser (150), His (229) | Nil |
| *Pseudomonas nanhaiensis* | Ser (101), Ser (171) | Phe (99), Val (218) |
| *Candidatus pseudomonas excrementarium* | 172 (Ser), Leu (217) | Ile (218) |
| *Acidovorax delafieldii* | Tyr (87), Thr (88) | Ile (222), His (251) |
| *Ketobacter alkanivorans* | Ser (108), Tyr (110), Ser (181), Trp (205) | Met (182), Val (229) |
| *Piscinibacter sp.* | Tyr (88), Ser (161), His (238) | Ile (209), His (238) |
| *Rhizobacter sp.* | Tyr (87) | Ile (208), His (237) |

Comparing the docking score between the predicted structures and with the one obtained for IsPETase allowed us to rate the proteins preliminarily as prospective PETases. While the best docking score obtained, as mentioned earlier, was −5.5 kcal/mol, some of the structures showed either low docking scores or had unacceptable values of RMSD. Of the 27 structures considered (including RgPETase), 15 structures showed binding affinity within −4.5 kcal/mol with acceptable RMSD values. Of these 15, 8 had docking scores very close to IsPETase, and 5 of these had identical docking pockets. (figures 2 and 3, table 2).

**Figure 3-.**
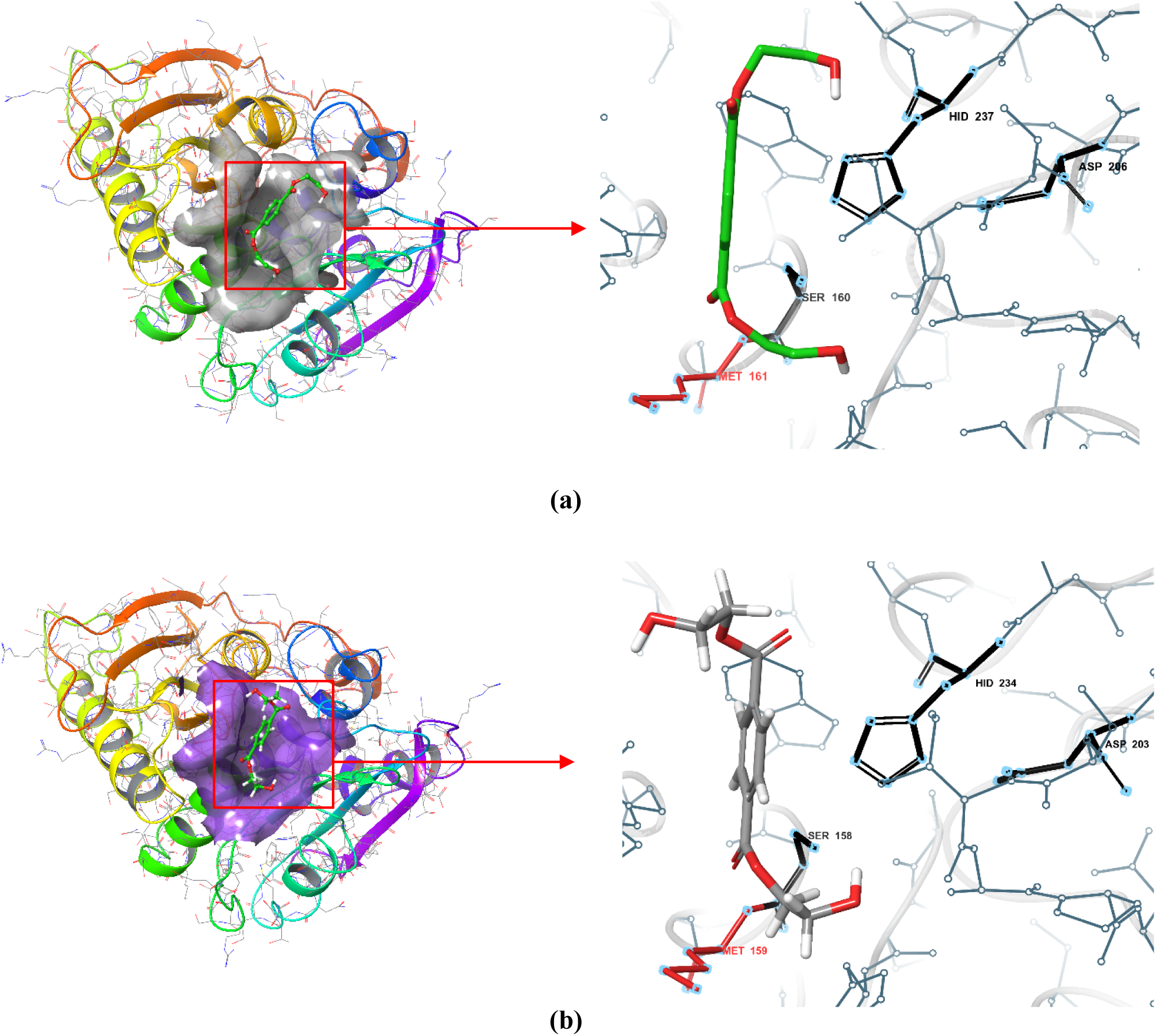
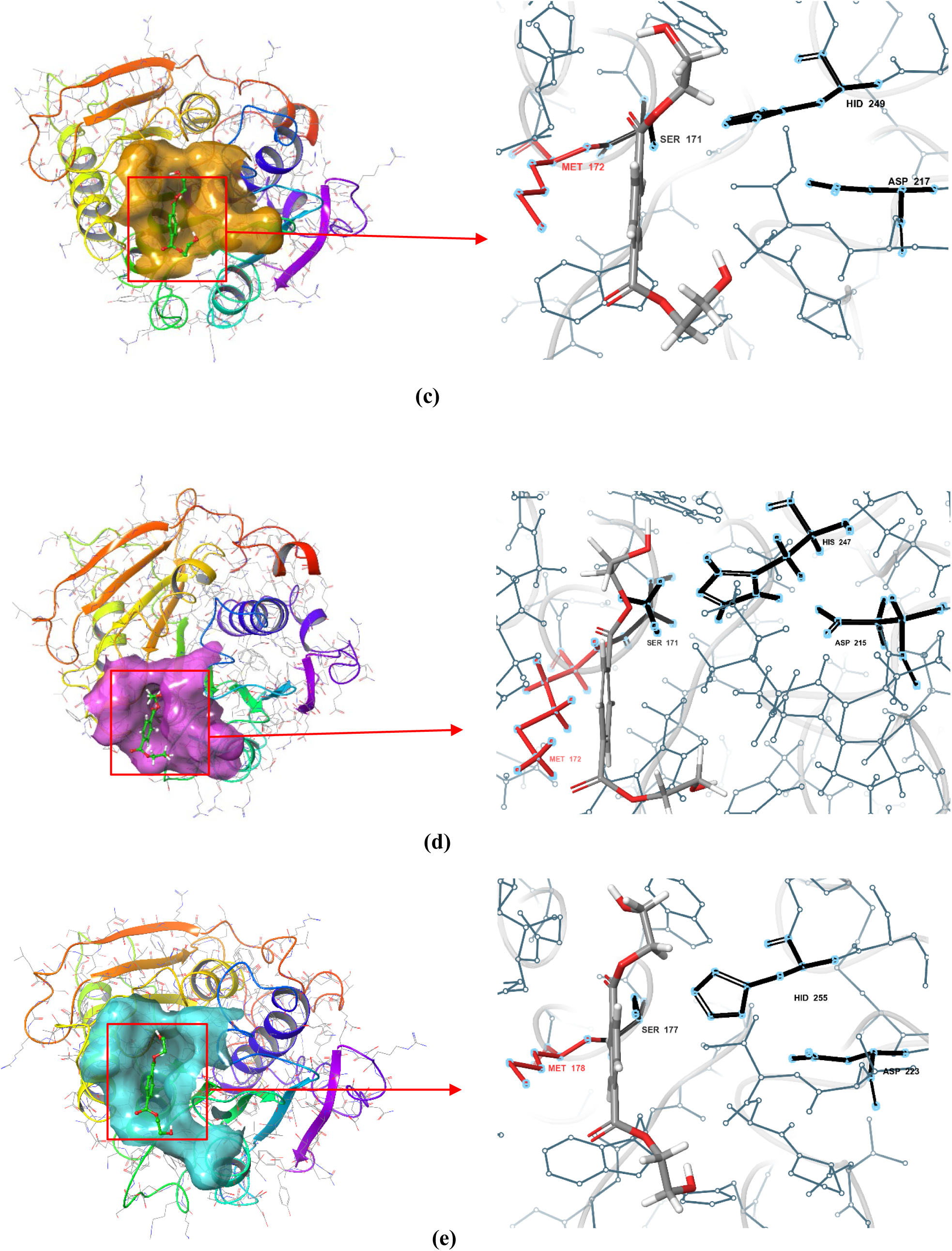
The docking pose illustrates the ligand’s accommodation within the protein’s binding pocket. This 3D visualisation elucidates the ligand’s orientation, location, and interactions with critical residues, offering insights into the binding mode and possible efficacy of the complex. The conserved catalytic residues are denoted in black, while Met is indicated in red in all the magnified images. a) PETase from *Ideonella sakaiensis* b) PETase from *Rhizobacter gummiphilus*. c) PETase from *Halopseudomonas pertucinogena* d) PETase from *Halopseudomonas bauzanensis* e) PETase from *Ketobacter sp*

**Table 2:** Docking score, Lower Bond (L.B.) RMSD Values and Upper Bond (U.B.) RMSD Values.

| No | Bacteria Name | Docking Score | L.B. RMSD Values | U.B. RMSD Values |
| --- | --- | --- | --- | --- |
| 1 | *Ideonella sakaiensis* | -5.1 | 1.183 | 1.702 |
| 2 | *Rhizobacter gummiphilus* | -5.4 | 1.244 | 0.505 |
| 3 | *Marinobacter daqiaonensis* | -5.5 | 1.372 | 1.041 |
| 4 | *Halopseudomonas pertucinogena* | -5.5 | 1.905 | 1.341 |
| 5 | *Pseudomonas zhaodongensis* | -5.3 | 1.157 | 0.202 |
| 6 | *Halopseudomonas bauzanensis* | -5.4 | 1.682 | 1.259 |
| 7 | *Ketobacter sp.* | -5.2 | 1.916 | 1.447 |
| 8 | *Burkholderia sp.* | -5.3 | 1.609 | 1.356 |
| 9 | *Aquabacterium sp.* | -5.1 | 1.434 | 0.932 |
| 10 | *Candidatus muproteobacteria bacterium* | -5.1 | 1.88 | 1.136 |
| 11 | *Pseudomonas nanhaiensis* | -5.1 | 2.01 | 1.215 |
| 12 | *Candidatus pseudomonas excrementarium* | -4.7 | 1.148 | 0.146 |
| 13 | *Acidovorax delafieldii* | -4.6 | 1.79 | 1.048 |
| 14 | *Ketobacter alkanivorans* | -4.6 | 1.665 | 1.183 |
| 15 | *Piscinibacter sp.* | -4.6 | 1.566 | 1.306 |
| 16 | *Rhizobacter sp.* | -4.5 | 1.412 | 1.105 |

HpPETase, HbPETase, and KsPETase belong to the type II class of PETases, exhibiting specific substitutions in the AXE domain and in subsite 1 and subsite 2, which influence the substrate binding position. The binding pockets of type II PETases exhibit modest variations and thereby, the binding conformation of the ligand differs from that of IsPETase, which is a type IIb class of PETase.

The serine, which was earlier mentioned as one of the conserved residues in the AXE domain, was expectedly present almost all the complexes formed, as either the donor or acceptor in the formation of hydrogen bonds with the BHET. Similarly, the presence of a tyrosine at position 87 of subsite 1 is deemed to favour an interaction between the aromatic group of the BHET and the PETase [30]. Among the predicted PETases, there is an equivalent distribution of tyrosine and phenylalanine at this position, which seems to be favourably interacting with the BHET as well. HpPETase, HbPETase and KsPETase showed the presence of a phenylalanine at this position. Traditionally, the nitrogen atoms of Y87 and M161 have been known to be involved in the formation of the oxyanion hole, found in type II PETases. In the case of HpPETase, HbPETase and KsPETase, the nitrogen from the phenylalanine and the methionine contribute to the oxyanion hole which stabilizes the negatively charged oxyanion intermediate during PET hydrolysis. The W185 in the subsite 1, however, is a residue which has been conserved across all the five PETases. Similarly, the isoleucine at 208, a part of subsite 1, has been found to be conserved in HpPETase and HbPETase, but has been replaced by a valine (hydrophobic) in RgPETase and KsPETase. Going by the activity of RgPETase, the substitution has minimal effect on the surface hydrophobicity.

In the subsite 2, W159 and N241 are highly conserved across all the PETases and has been found to be crucial for the enzymatic function, while A89 is replaced by a serine in RgPETase, HpPETase, HbPETase and KsPETase. Despite the substitution, there has been no reduction in the hydrolase activity of RgPETase when compared to IsPETase [14]. The threonine at position 88 is however a determinant in the classification of type II PETases into subcategories a and b. While T88 is highly conserved in type IIb PETases [6], substitutions of threonine to valine or leucine has been noted in type IIa PETases. The presence of a T88L mutation in RgPETase and T88V in KsPETase, HpPETase, and HbPETase, are indicative of their classification as type IIa PETases [6]. Similarly, Ser238 is substituted by phenylalanine in RgPETase and KsPETase, but in HpPETase and HbPETase, it is substituted by a tyrosine, which further point to their type IIa categorisation. All these substitutions have significantly improved the binding of substrate, with HpPETase showing four times higher binding free energy, while KsPETase showed two times higher binding free energy when compared with the IsPETase. When compared with RgPETase (also a type IIa PETase), all three candidates have substitutions in subsite II and show better binding free energy.

The docking pose related to the lowest docking scores were selected for further MD simulations. The pose was manually checked for the location, orientation and interactions of the ligand with the key binding site residues and the absence of unfavorable bonds. The docking poses chosen for the MD were therefore based on the correctness of the orientation and not the mechanical choice of the topmost poses (table 2).

The binding free energy of protein-ligand complexes was calculated using MMPBSA to assess their stability and binding affinity. The cumulative binding free energies for the five prospective PETase-BHET (IsPETase-BHET, RgPETase-BHET, HpPETase-BHET, HbPETase-BHET, and KsPETase-BHET) complexes were −2.75 ± 3.41, −0.59 ± 3.45, −12.16 ± 3.02, −1.98 ± 3.93, and −6.14 ± 3.14 kcal/mol, respectively, as presented in table 3. The PETase from *Halopseudomonas pertucinogena* has the most robust binding relative to the other PETases in the list, having nearly four-fold binding affinity than IsPETase, whereas, the PETase derived from *Ketobacter sp.* exhibits twice the binding affinity in comparison to IsPETase. PETase from *Halopseudomonas bauzanensis* exhibits somewhat lower binding free energy than IsPETase, while RgPETase had the lowest binding affinity for BHET.

**Table 3:** MMPBSA Binding Free energy of protein-ligand complex.

| **PETase** | **MMPBSA Binding Free Energy ($\Delta G$ binding) (Kcal/mol)** |
| --- | --- |
| *Ideonella sakaiensis* | -2.75 +/- 3.41 |
| *Rhizobacter gummiphilus* | -0.59 +/- 3.45 |
| *Halopseudomonas pertucinogena* | -12.16 +/- 3.02 |
| *Halopseudomonas bauzanensis* | -1.98 +/- 3.93 |
| *Ketobacter sp.* | -6.14 +/- 3.14 |

Solvation being a crucial factor, influencing the protein’s folding and stability, the SASA values for the PETases were also looked at showing the least solvent-accessible surface area for *I. sakaiensis* and *R. gummiphilus* (figure 4 a,b). The other PETases – from *H. pertucinogena* (HpPETase)*, H. bauzanensis* (HbPETase) and *Ketobacter* sp. (KsPETase) also had comparable SASA values (figure 4c, 4d and 4e). Among the surface amino acids contributing to the binding, a methionine near the serine hydrolase motif in the AXE domain was found to be omnipresent and almost always had a SASA value of zero in the ligand-bound complex, indicative of its role in binding BHET (figure 3). Likewise, this methionine was also found to be involved in the hydrophobic interactions between the PETase and its target.

**Figure 4-.**
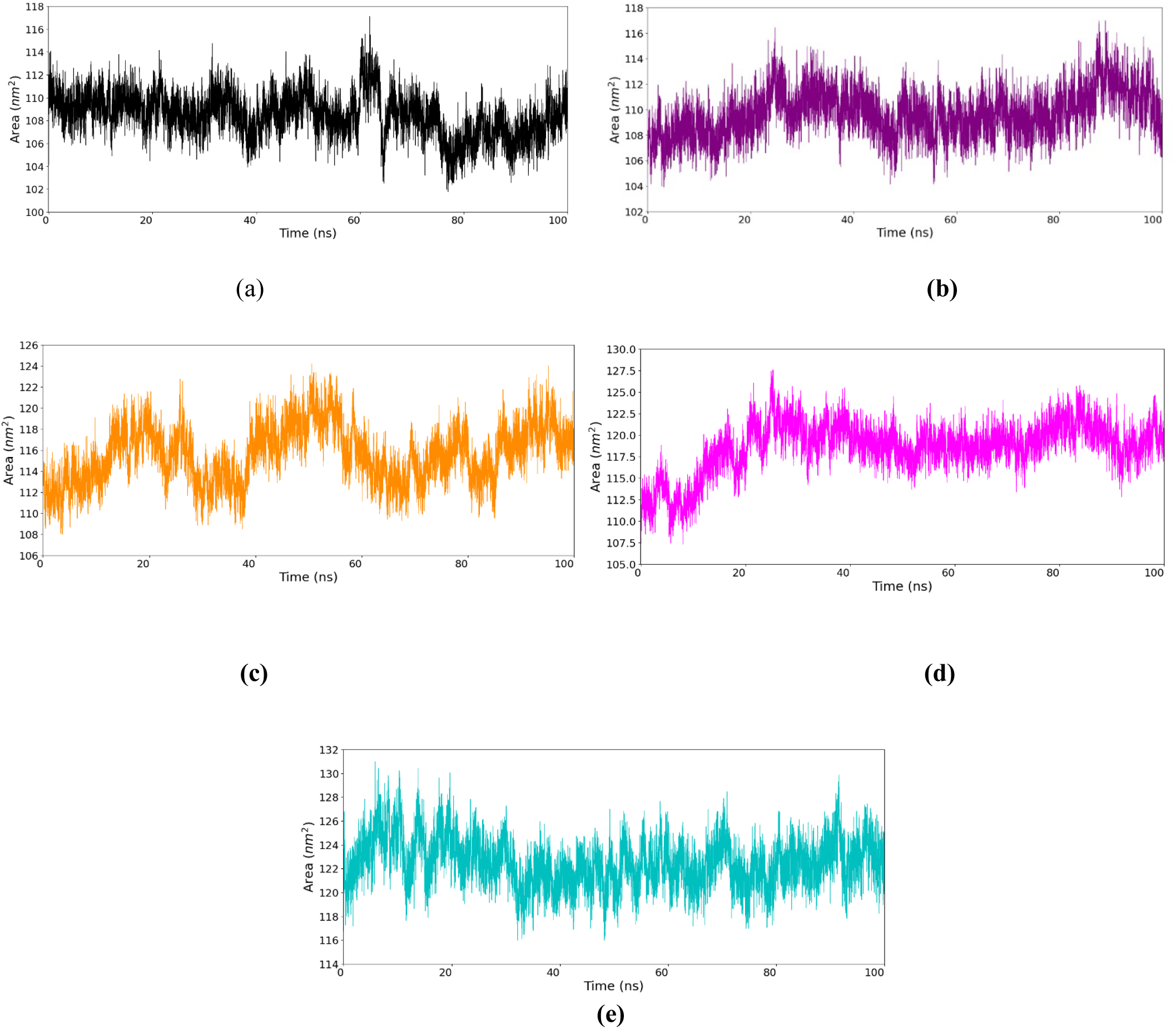
The Solvent Accessible Surface Area (SASA) plot for the protein-ligand complex during the molecular dynamics simulation illustrates the degree of the protein’s surface exposure to the solvent over time. Variations in SASA values indicate conformational changes, folding processes, or modifications in the stability of the complex. a) PETase from *Ideonella sakaiensis* (black colour) b) PETase from *Rhizobacter gummiphilus* (purple) c) PETase from *Halopseudomonas pertucinogena* (orange) d) PETase from *Halopseudomonas bauzanensis* (pink) e) PETase from *Ketobacter sp.* (cyan)

### Stability of PETase-BHET interactions

The detailed secondary structural transitions of PETase in presence of BHET were studied employing MD simulations for 100 ns. In order to mitigate the potential for poor connections and geometry inside the structure, energy minimization was conducted prior to conducting the molecular dynamics (MD) simulation.

Among all the PETase enzymes, HpPETase was found to be the most resilient on interaction with BHET (even better than IsPETase), with the RMSD values stabilizing around 0.4 nm throughout the simulation, after an initial inflection (figure 5c). In case of the IsPETase-BHET complex, the average root mean square deviation (RMSD) values remained consistently below 0.5 nm for the whole simulation. The HbPETase-BHET complex exhibited the third highest stability, with an average RMSD value of 0.6 nm. The KsPETase performed better than the RgPETase in it’s interactions with BHET – the RMSD stabilizing around 0.8 nm in due course, albeit with intermediate flexions. The RgPETase, which has now been experimentally proven to degrade PET, stabilized at around a RMSD value of 1 nm. Overall, the dynamics for all the 5 PETases suggested that the structures of the proteins remained intact during the simulation (figure 5). In each RMSD plot, the darker lines highlight the RMSD during the protein-ligand interaction against the RMSD of the protein backbone (in faded lines).

**Figure 5:**
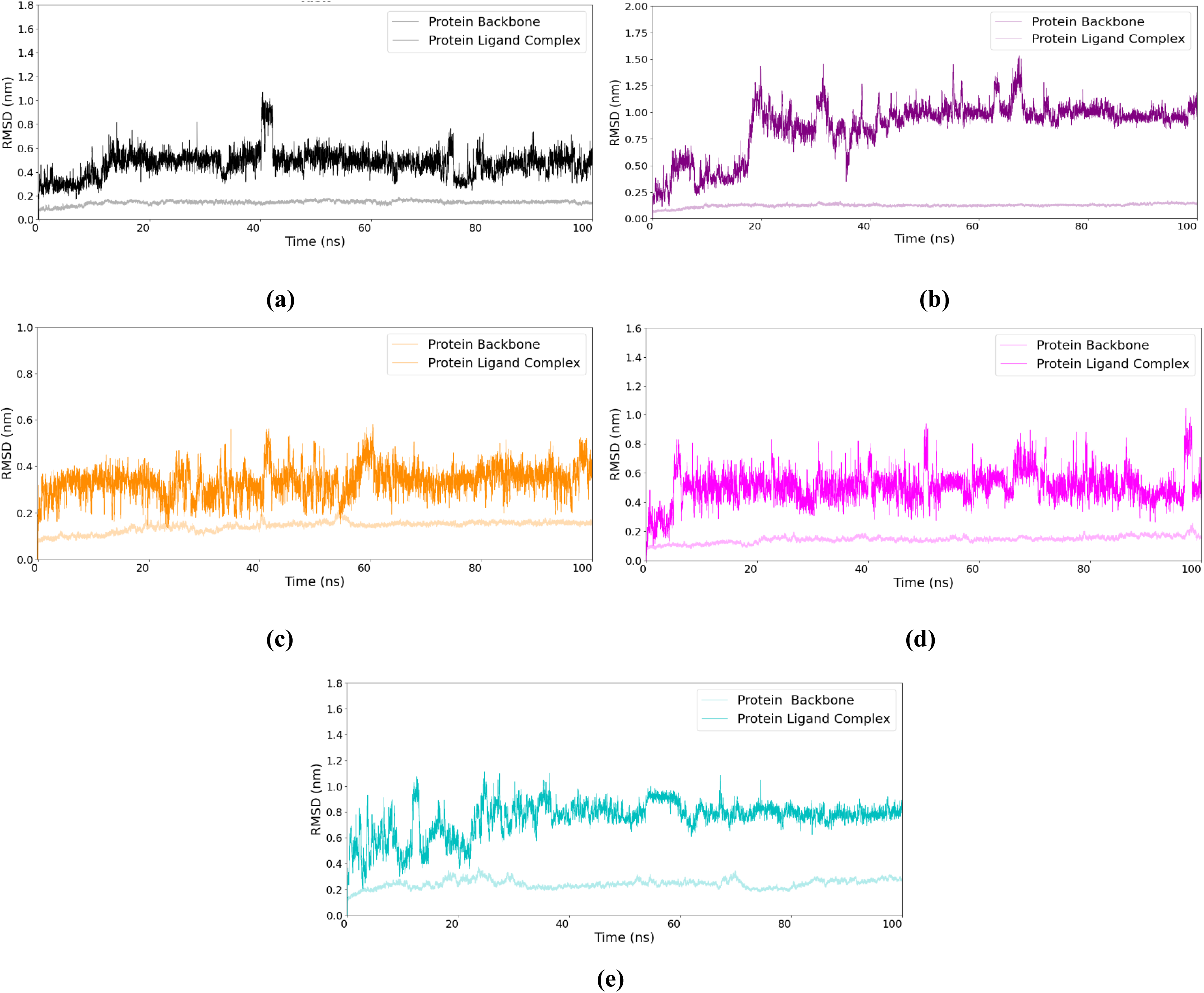
The RMSD plots demonstrate the structural stability of the protein-ligand complex throughout the duration of simulation. Consistent RMSD values indicate that the complex has reached equilibrium, signifying stable interactions between the protein and ligand. In each figure bright colour lines represent protein-ligand RMSD and light colour lines are representing the protein backbone a) PETase from *Ideonella sakaiensis* (black colour) b) PETase from *Rhizobacter gummiphilus* (purple colour) c) PETase from *Halopseudomonas pertucinogena* (orange colour) d) PETase from *Halopseudomonas bauzanensis* (pink colour) e) PETase from *Ketobacter sp.* (cyan colour)

The root mean square fluctuation (RMSF) was calculated to analyze the mobility and flexibility of individual amino acids from their mean positions during simulation, which allows the identification of the specific areas of the protein showing higher degrees of dynamic behavior. The RMSF values consistently stayed below 0.3 for all the PETases, with occasional peaks above 0.3 (+0.1) (figure 6). In all the figures, the curve for RMSF of the specific PETase has been contrasted with the curve for IsPETase (shown in the background in grey). For the reported PETases of *I. sakaiensis* and *R. gummiphilus*, the RMSF values were identical and lower in comparison to the other PETases. Alignment of the sequences of the PETases with the RMSF data revealed that the fluctuations were greater in the regions of the proteins having flexible loops, as compared to the more structured regions (supplementary table 5). The N terminal residues which show increased flexibility are predictably the ones with greater exposure to the solvent. The peaks of significant intensity visible in the RMSF plot of HbPETase, KsPETase, and HpPETase (figure 6b, 6c and 6d) are indicative of the presence of significantly more flexible regions of the protein segment at multiple places in these enzymes. The substitution and addition events observed in the sequence alignment have an impact on the peak heights observed in RMSF. Though the loop regions display increased variability in these PETases, this has however not impacted the ligand binding, as none of the residues from these regions contributed to the ligand binding cavity. The residues comprising the binding site of all the proteins exhibited a high degree of stability and fluctuations, if any were of relatively insignificant magnitude. The orange colour used in the figure indicates regions of intense peaks, while specific information relating to the residues can be found in the supplementary table 5.

**Figure 6:**
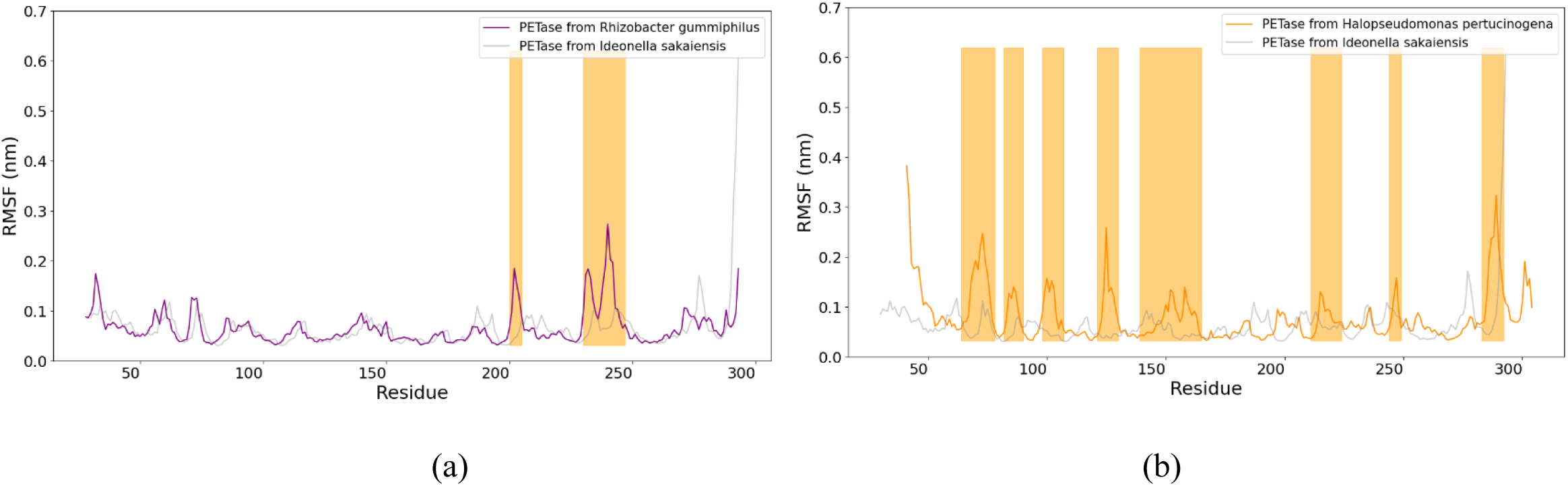
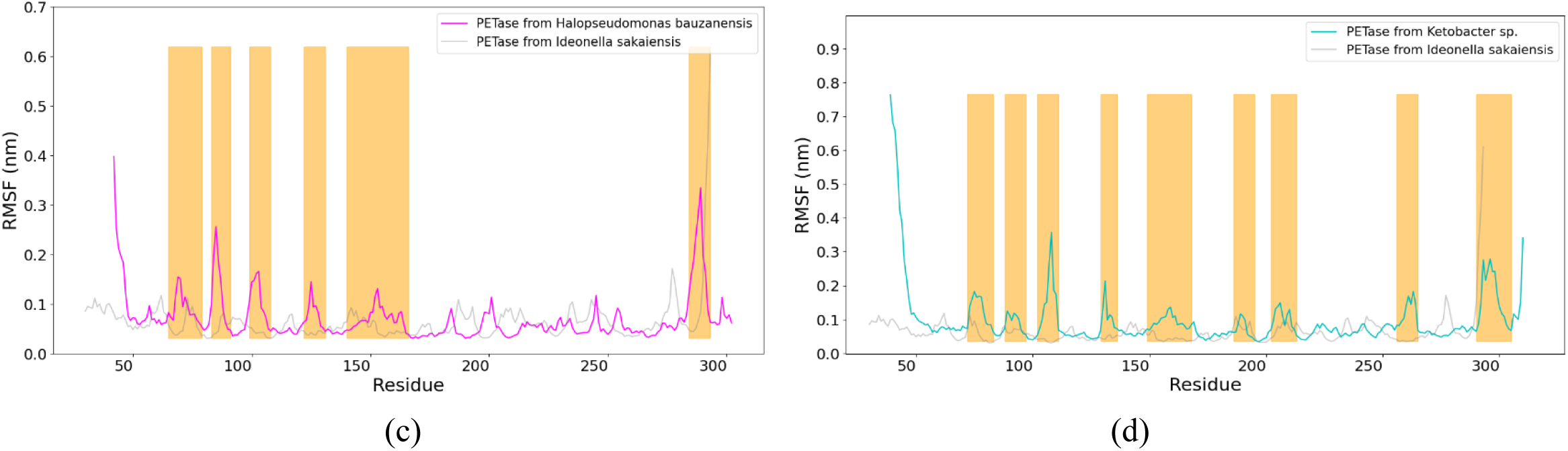
The Root Mean Square Fluctuation (RMSF) plot for the protein-ligand complex during the molecular dynamics simulation demonstrates the temporal flexibility of each residue. Peaks in the RMSF profile signify places of increased mobility, such as loops or terminal ends, whereas valleys denote more stable, stiff areas. In each plot, the RMSF of protein residues were compared with IsPETase. The RMSF of IsPETase is depicted in grey in each plot. a) PETase from *Rhizobacter gummiphilus*. b) PETase from *Halopseudomonas pertucinogena* c) PETase from *Halopseudomonas bauzanensis* d) PETase from *Ketobacter sp*.

These results are also in correlation with the Free Energy Landscape plots (figure 7). While the free energy minima for IsPETase (figure 7a) and RgPETase (figure 7b) are quite similar, HpPEtase, HbPETase, and KsPETase enzymes too exhibit suitably low free energy levels in the presence of the BHET ligand (figure 7c, 7d and 7e), which indicates structural stability throughout the simulation. The radius of gyration of the five PETases were also within permissible range showing acceptable stability and maintenance of conformational compaction during the simulation (figure 8). However, the radius of gyration showed more fluctuations in the KsPETase (figure 8e), indicating a higher order of flexibility.

**Figure 7:**
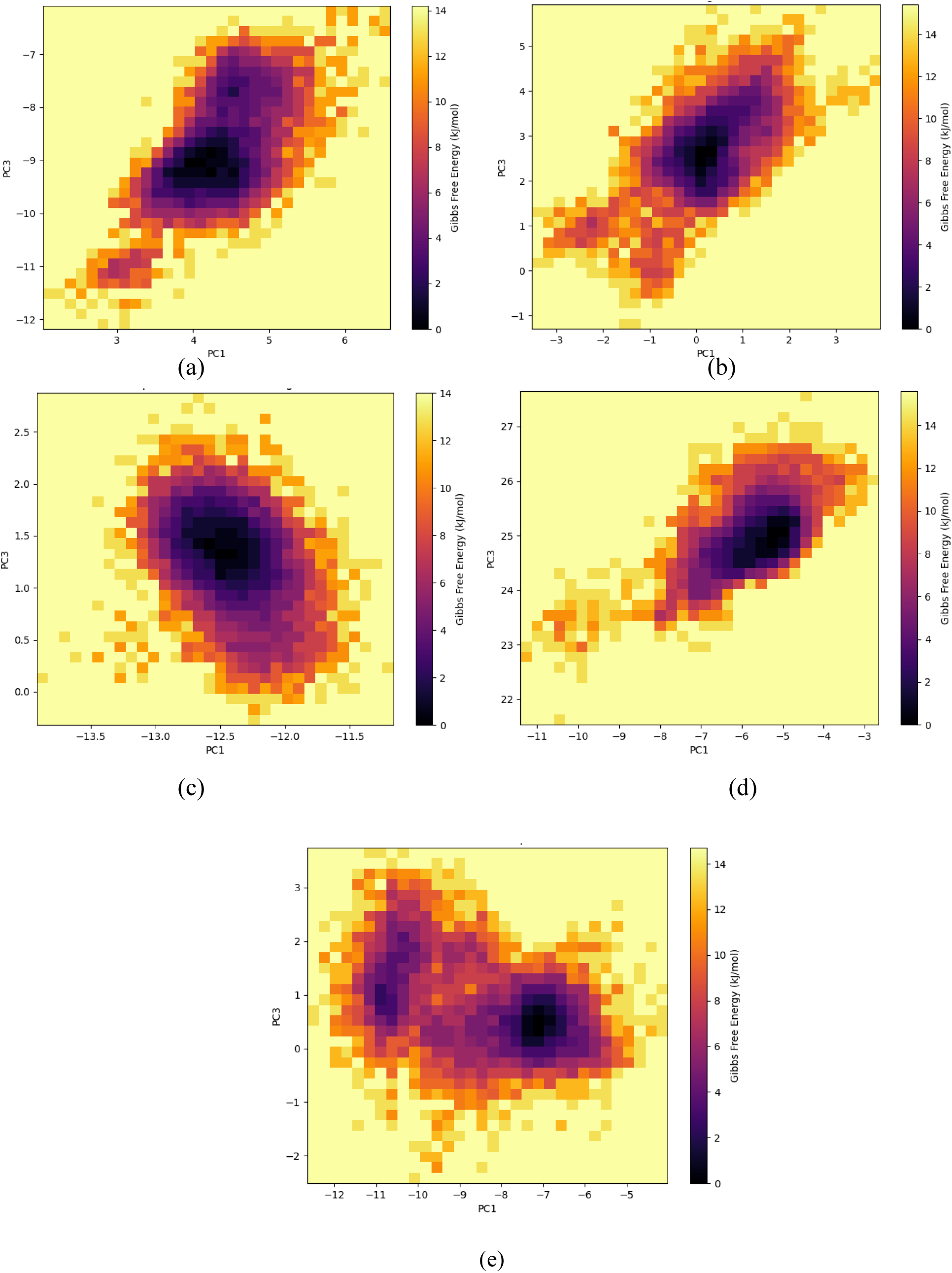
Free energy landscape between PC1 and PC3 of the different PETases with BHET (a) IsPETase (b) RgPETase (c) HpPETase (d) HbPETase (e) KsPETase

**Figure 8:**
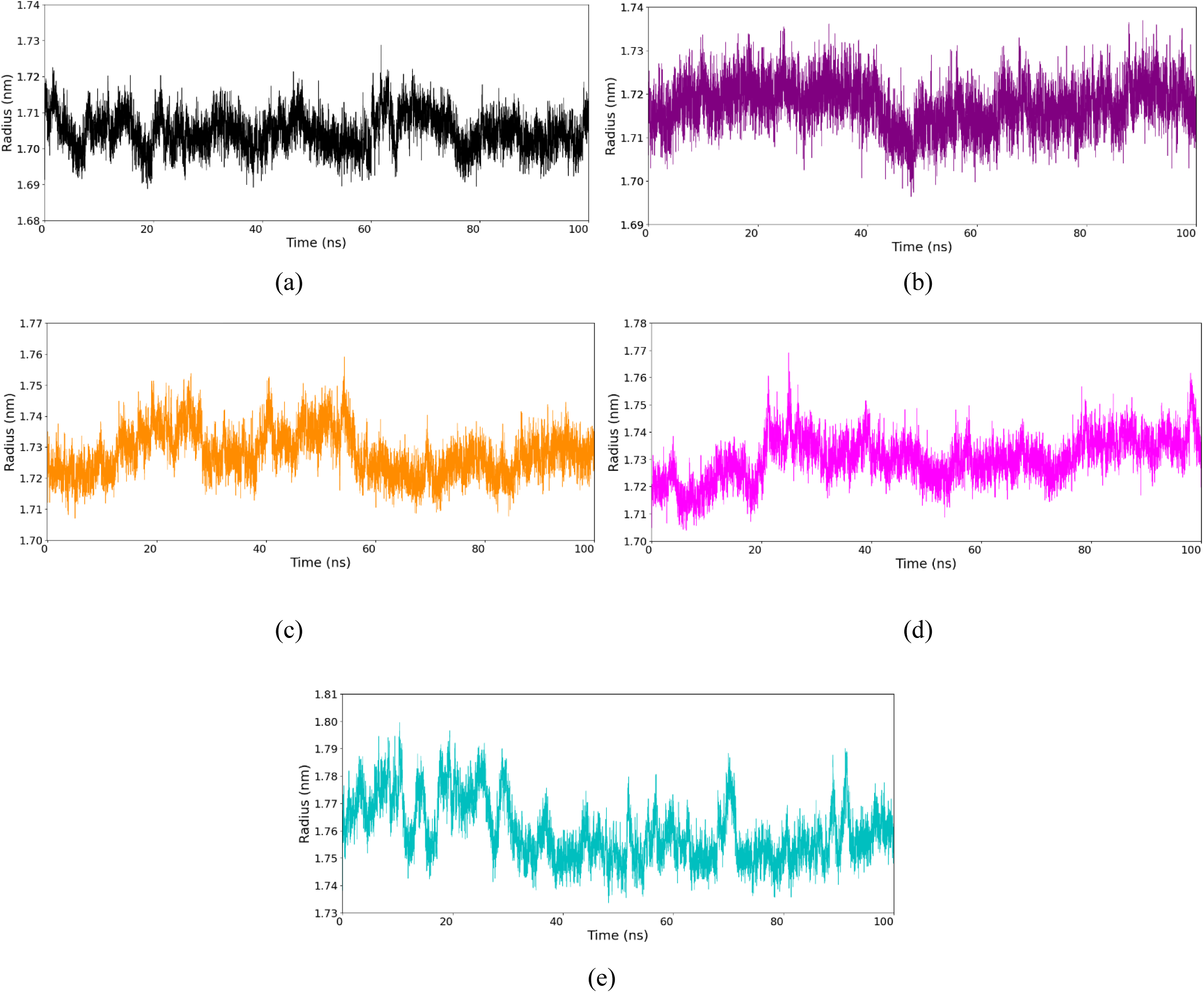
The Radius of Gyration (Rg) plot for the protein-ligand complex during the molecular dynamics simulation measures the compactness of the complex over time. Stable Rg values suggest that the complex maintains its folded structure throughout the simulation, indicating consistent interactions between the protein and ligand. 1) PETase from *Ideonella sakaiensis* (black colour) 2) PETase from *Rhizobacter gummiphilus* (purple) 3) PETase from *Halopseudomonas pertucinogena* (orange) 4) PETase from *Halopseudomonas bauzanensis* (pink) 5) PETase from *Ketobacter sp.* (cyan)

Principal Component Analysis was performed to identify collective motion of the different PETases. Quantitative understanding of the directions of motion captured by the eigenvectors were depicted the 2D projection of eigen vectors 1 and 3 (principal components 1 and 3) on the resulting 100 nanosecond molecular dynamics trajectories. In order to simplify the calculation, the covariance matrix of Cα atomic fluctuations was converted into a diagonal matrix to determine the eigen values. A denser and more localized area in the conformational space corresponds to a stable ensemble, with very few structural variations. On the other hand, a more dispersed distribution with multiple clusters would be representative of greater conformational flexibility. The PCA scatter plots of IsPETase, HpPETase and HbPETase (figures 9a, c and d) thereby reveal a singular predominant conformation with no significant dispersion during the simulation process. On the other hand, the plot for RgPETase (figure 9b), which shoes more diversity (two distinct clusters) is reflective of a transition between states, resulting mostly from the ligand binding. The plot for KsPETase shows an even larger diffusiveness (more than two clusters), indicating a transition between at least two different conformational states during the interaction with the ligand. The observations reported for all the PETases captured more than 99% of the total motion, and hence, reveal a complete landscape of the interactions.

**Figure 9:**
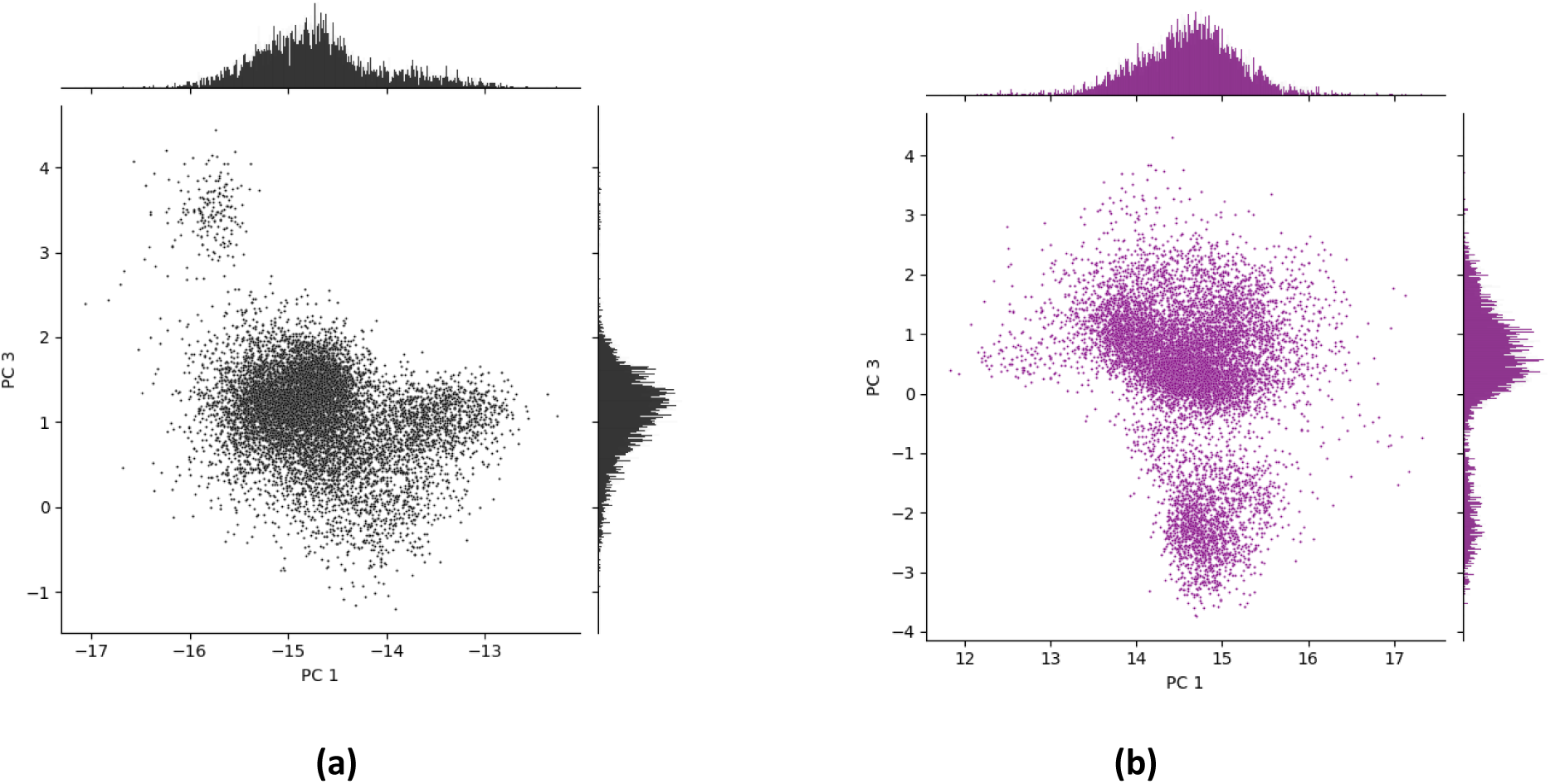
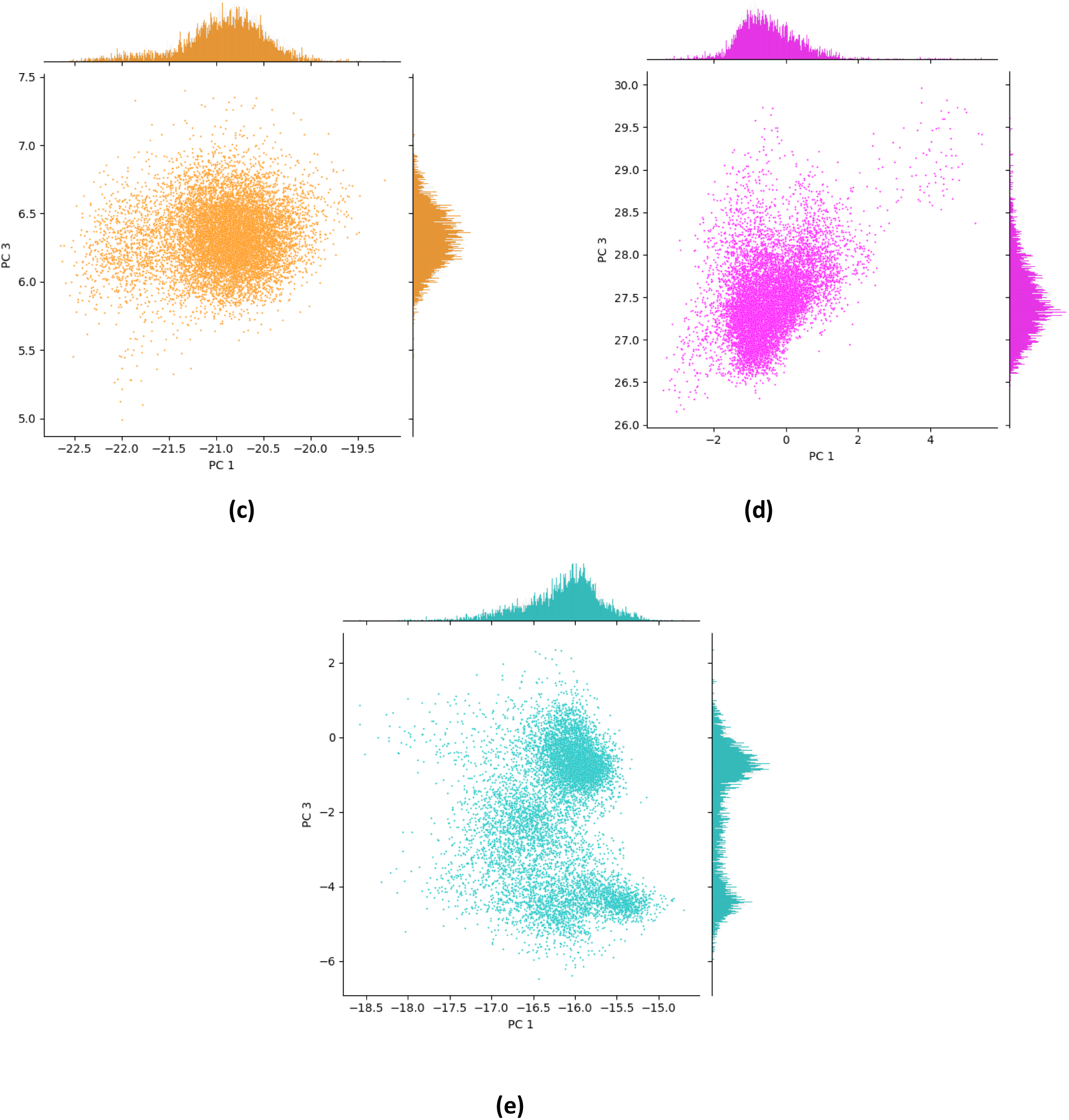
The PCA scatter plots of PC1 vs PC3 illustrate the principal patterns of motion in the protein-ligand complex throughout the simulation. The clear grouping or dispersion in the figure signifies the extent of conformational alterations and dynamic behaviour of the complex. a) PETase from *Ideonella sakaiensis* (black colour) b) PETase from *Rhizobacter gummiphilus* (purple colour) c) PETase from *Halopseudomonas pertucinogena* (orange colour) d) PETase from *Halopseudomonas bauzanensis* (pink colour) e) PETase from *Ketobacter sp.* (cyan colour)

## Conclusion

The discovery of a PETase from *I. sakaiensis* with high degradability and specificity in the recent past provided a major boost to the search for plastic degrading enzymes from microbes. The subsequent discovery of *R. gummiphilus* stirred the interest even more, and the past few years have seen renewed vigour in the quest for a naturally occurring solution to the ever-increasing threat of plastic pollution in the environment. The work in this manuscript would supplant and provide a headstart to these efforts as it reveals a handful of prospective bacterial candidates which could be tapped to harness their plastic degradation abilities.

All of the species shortlisted for this work have shown the presence of a conserved hydrolysis domain, and some of them have been documented to hydrolyze PET and other polymers. Substitution in the catalytic triad has been demonstrated to diminish enzyme activity, and sequence alignment confirms that the catalytic triad residues are 100% conserved across all enzymes. Besides the typified Ser160, Asp206 and His 237 catalytic triad, the presence of a conserved Met161 in the vicinity has also been revealed as a major contributor to binding targets. All the three candidates have substitutions akin to the type IIa category of PETases, further bolstered by the two conserved cysteine residues at positions 203 and 239 for the formation of an extra disulphide bond (unlike type I PETases).

The spectrum of variation in the source of these bacterial candidates is also diverse and includes species residing in environments ranging from the marine to sub-zero habitats. *Ketobacter alkanivorans* [31], *Marinobacter daqiaonensis* [32], and *Gammaproteobacteria* bacterium [33] were identified from marine sources and possess conserved hydrolase domains that may aid in the breakdown of microplastics in the ocean, a major issue that plagues the environment and mankind. On the other hand, *Oleispira antarctica* is another bacterium that could be instrumental in plastic degradation at lower temperatures[34]. *A.delafieldii* is a common Gram negative aerobe which had previously exhibited high degradation activity against Poly(tetramethylene succinate)-co-(tetramethylene adipate) (PBSA) [35] are commonly found in soil, water and on plants but are seldom implicated in human infections Interestingly, *Schlegelella brevitalea* has been identified as a triple mutant of *Ideonella sakaiensis*, although its hydrolysis activity may not be as efficient as that of *I. sakaiensis* [36]. The species of *Pseudomonas* that have been reported, are known to survive across variable environmental conditions and can prospectively degrade plastics to varying degrees [37].

The notable conservation of the AXE domain among all the examined PETases, particularly the essential residues in the binding locations (including subsite1 and subsite 2), strongly indicates an evolutionary inclination for plastic degradation abilities. The MD studies of the five PETases from various species revealed significant potential in HpPETase, HbPETase, and KsPETase for plastic degradation, comparable to the experimentally validated IsPETase and RgPETase. The binding of BHET to HpPETase, infact, showed better binding energies than the reference IsPETase and RgPETase, signifying improved substrate affinity. The values of the MMPBSA calculations in conjunction with its RMSD plot, illustrate enhanced structural stability over time for HpPETase-BHET binding, when contrasted with both IsPETase and RgPETase. The same could also be hypothesized for the HbPETase and the KsPETase too, arguably to a lesser extent though. The conformational stability of the PETases during the simulation using the PCA analysis again revealed stable structural conformations for the HpPETase, comparable to IsPETase. Interestingly, the PETase from *R. gummiphilus* (one of the validated PETases) displayed a rather diffusive trajectory during the dynamics, suggesting a flexible conformation during binding interactions, which may be attributed to the presence of variable region 1 (VR1, Q26-S51), the wobbling tryptophan-containing loop (WW-loop, W183-S187), variable region 2 (VR2, P207-V228) and the C-terminal region (G271-Y292) in the structure of this PETase. Furthermore, the outputs of Free Energy Landscape (FEL) and Radius of Gyration (Rg) consistently demonstrated that these potential PETases preserve structural integrity and compactness during the simulation duration.

The PETase from *H. pertucinogena* and to some extent, *H. bauzanensis* can be therefore, subjected to experimental analysis of their plastic degradation activity. Indeed, a structure for the PETase from *H. bauzanensis* have been now deposited, but the reports of its activity still remains elusive. The prospective PETase from *Ketobacter sp.* however, does not show as much promise as the other two, but in some aspects, resembles the binding interactions of *R. gummiphilus* and hence, can also be tried for PET degradation. Although the results obtained in the current study suggest the PET degradation activities of these three alone, the rest of the PETases could also be a source of plastic degradation enzymes, with suitable protein engineering processes to increase their activity. In that direction, the characterization of the interface in these prospective enzymes, with a detailed understanding of the surface amino acids and their binding mechanisms is also supposed to provide a significant platform to the process.

## Declarations

### Ethics approval and consent to participate

Not Applicable

### Consent for publication

All the authors have read the manuscript and have provided consent for publication.

### Availability of data and materials

The data supporting the results are provided with the supplementary files and is open for access to readers.

### Authors’ contributions

YR has been responsible for the acquisition and analysis of the data, investigation, prepared figures and tables and also scripted the first draft of the manuscript. SB has been involved in the ideation, analysis, validation of data and the final version of the manuscript.

## Acknowledgements

YR has been supported by a fellowship from BITS, Pilani during the tenure of this work.

