## supplementary for "Identification of Prospective PETases Across Prokaryotes Using an *in silico* Approach"

**Supplementary Table 1:** Blast output against the IsPETase sequence.

| Accession | Scientific name | Confidence (%) | Sequence identity (%) | Alignment coverage (%) | Hit info |
|---|---|---|---|---|---|
| WP_223655417.1 | Pseudomonas_nanhaiensis | 100 | 75 | 85 | Chain: A: PDB Molecule:polyester hydrolase; |
| BAB86909.1 | Acidovorax_delafieldii | 100 | 81 | 85 | Chain: A: PDB Molecule:poly(ethylene terephthalate) hydrolase; |
| WP_036989706.1 | Halopseudomonas_bauzanensis | 100 | 81 | 86 | Chain: A: PDB Molecule:polyester hydrolase; |
| WP_122164848.1 | Pseudomonas_zhaodongensis | 100 | 63 | 90 | Chain: A: PDB Molecule:cutinase; |
| WP_092009807.1 | Marinobacter_daqiaonensis | 100 | 75 | 84 | Chain: A: PDB Molecule:polyester hydrolase; |
| WP_185266504.1 | Halopseudomonas_xiamenensis | 100 | 79 | 86 | Chain: A: PDB Molecule:polyester hydrolase; |
| WP_156871850.1 | Ketobacter_sp._MCCC_1A13808 | 100 | 65 | 87 | Chain: A: PDB Molecule:cutinase; |
| WP_090272969.1 | Halopseudomonas_litoralis | 100 | 80 | 86 | Chain: A: PDB Molecule:polyester hydrolase; |
| WP_047194864.1 | Schlegelella_brevitalea | 100 | 70 | 89 | Chain: B: PDB Molecule:dlh domain- |

| | | | | | |
|---|---|---|---|---|---|
| | | | | | containing protein; |
| MBU1194958.1 | Proteobacteria_bacterium | 100 | 56 | 61 | Chain: B: PDB Molecule:dlh domain-containing protein; |
| OGI66810.1 | Candidatus_Muproteobacteria_bacterium_RBG_16_60_9 | 100 | 69 | 93 | Chain: B: PDB Molecule:dlh domain-containing protein; |
| MBC7701646.1 | Massilia_sp.7 | 100 | 57 | 89 | Chain: B: PDB Molecule:dlh domain-containing protein; |
| MBC7956640.1 | Cytophagales_bacterium | 100 | 85 | 90 | Chain: A: PDB Molecule:poly(ethylene terephthalate) hydrolase; |
| WP_175819463.1 | Burkholderia_sp._BCC0419 | 100 | 57 | 85 | Chain: B: PDB Molecule:dlh domain-containing protein; |
| WP_054022242.1 | Piscinibacter_sakaiensis | 100 | 100 | 100 | Chain: A: PDB Molecule:poly(ethylene terephthalate) hydrolase; |
| WP_090538641.1 | Halopseudomonas_formosensis | 100 | 83 | 86 | Chain: A: PDB Molecule:polyester hydrolase; |
| HIZ51049.1 | Candidatus_Pseudomonas_excrementavium | 100 | 83 | 86 | Chain: A: PDB Molecule:polyester hydrolase; |
| PKM05449.1 | Gammaproteobacteria_bacterium_HGW-Gammaproteobacteria-6 | 100 | 80 | 84 | Chain: A: PDB |

| | | | | | |
|---|---|---|---|---|---|
| | | | | | Molecule:polyester hydrolase; |
| WP_085749238.1 | Rhizobacter_gummiphilus | 100 | 100 | 91 | Chain: A: PDB Molecule:rgcutii; |
| WP_220809760.1 | Noviherbaspirillum_aridicola | 100 | 75 | 59 | Chain: B: PDB Molecule:dlh domain-containing protein; |
| WP_101893885.1 | Ketobacter_alkanivorans | 100 | 66 | 84 | Chain: A: PDB Molecule:cutinase; |
| MBQ0729274.1 | Oleispira_antarctica | 100 | 60 | 84 | Chain: A: PDB Molecule:lipiaf5-2; |
| WP_188635463.1 | Halopseudomonas_pertucinogena | 100 | 82 | 86 | Chain: A: PDB Molecule:polyester hydrolase; |
| MBX3625601.1 | Rhizobacter_sp. | 100 | 97 | 90 | Chain: A: PDB Molecule:poly(ethylene terephthalate) hydrolase; |
| MBA1146895.1 | Ectothiorhodospiraceae_bacterium_WFHF3C12 | 100 | 76 | 83 | Chain: A: PDB Molecule:lipiaf5-2; |
| MBI3384080.1 | Aquabacterium_sp. | 100 | 58 | 90 | Chain: B: PDB Molecule:dlh domain-containing protein; |
| MBL4611384.1 | Pseudomonas_sp. | 100 | 75 | 88 | Chain: A: PDB Molecule:polyester hydrolase; |

**Supplementary Table 2: Mutation analysis of AXE domain:** Substitution, addition, and deletion events in the AXE domain.

| PETase source organism | Substitution | Addition | Deletion | Total |
|---|---|---|---|---|
| *Acidovorax delafieldii* | 4 | 0 | 0 | 4 |
| *Ideonella sakaiensis* | 0 | 0 | 0 | 0 |
| *Candidatus Pseudomonas excrementavium* | 13 | 0 | 1 | 14 |
| *Ectothiorhodospiraceae bacterium* | 14 | 0 | 0 | 14 |
| *Massilia sp.* | 12 | 0 | 0 | 12 |
| *Cytophagales bacterium* | 4 | 0 | 0 | 4 |
| *Aquabacterium sp.* | 11 | 0 | 0 | 11 |
| *Pseudomonas sp.* | 10 | 0 | 1 | 11 |
| *Oleispira antarctica* | 12 | 0 | 1 | 13 |
| *Proteobacteria bacterium* | 12 | 1 | 0 | 13 |
| *Rhizobacter gummiphilus* | 0 | 0 | 0 | 0 |
| *Candidatus Muproteobacteria bacterium* | 7 | 1 | 0 | 8 |
| *Gammaproteobacteria bacterium HGW-Gammaproteobacteria-6* | 13 | 0 | 1 | 14 |
| *Halopseudomonas bauzanensis* | 13 | 0 | 1 | 14 |
| *Schlegelella brevitalea* | 8 | 0 | 0 | 8 |
| *Rhizobacter sp.* | 6 | 0 | 0 | 6 |
| *Halopseudomonas litoralis* | 13 | 0 | 1 | 14 |
| *Halopseudomonas formosensis* | 13 | 0 | 1 | 14 |
| *Marinobacter daqiaonensis* | 11 | 0 | 1 | 12 |
| *Ketobacter alkanivorans* | 12 | 0 | 1 | 13 |
| *Pseudomonas zhaodongensis* | 14 | 0 | 1 | 15 |
| *Ketobacter sp.* | 13 | 0 | 1 | 14 |
| *Burkholderia sp.* | 13 | 0 | 0 | 13 |
| *Halopseudomonas xiamenensis* | 13 | 0 | 1 | 14 |
| *Halopseudomonas pertucinogena* | 13 | 0 | 1 | 14 |
| *Piscinibacter sp. HJYY11* | 6 | 0 | 0 | 6 |
| *Noviherbaspirillum aridicola* | 10 | 0 | 0 | 10 |
| *Pseudomonas nanhaiensis* | 13 | 0 | 1 | 14 |
| Total | 283 | 2 | 14 | |

**Supplementary Table 3: Structure validation report:** The protein structures predicted using homology modelling were verified using PROCHECK and ERRAT. PROCHECK evaluated stereochemical quality by Ramachandran plot analysis, whereas ERRAT rated overall model reliability based on non-bonded interactions.

| No | Bacteria name | ProCheck (Residues in allowed region of Ramachandran plot) | ERRAT (Overall quality factor) |
|---|---|---|---|
| 1 | *Acidovorax delafieldii* | 100 % | 88.58 |
| 2 | *Candidatus Pseudomonas excrementavium* | 99.5% | 78.74 |
| 3 | *Ectothiorhodospiraceae bacterium* | 100% | 82.81 |
| 4 | *Massilia sp.* | 99.6% | 76.95 |
| 5 | *Cytophagales bacterium* | 100% | 89.02 |
| 6 | *Aquabacterium sp.* | 99.1% | 95.31 |
| 7 | *Pseudomonas sp.* | 98.6% | 73.33 |
| 8 | *Oleispira antarctica* | 100% | 78.43 |
| 9 | *Proteobacteria bacterium* | 99.1% | 62.93 |
| 10 | *Rhizobacter sp.* | 100% | 86.66 |
| 11 | *Candidatus Muproteobacteria bacterium* | 99.5% | 74.14 |
| 12 | *Gammaproteobacteria bacterium HGW-Gammaproteobacteria-6* | 99.1% | 68.37 |
| 13 | *Halopseudomonas bauzanensis* | 99.1% | 67.84 |
| 14 | *Schlegelella brevitalea* | 99.6% | 74.70 |
| 15 | *Rhizobacter gummiphilus* | 100% | 82.18 |
| 16 | *Halopseudomonas litoralis* | 99.1% | 70.58 |
| 17 | *Halopseudomonas formosensis* | 99.5% | 79.21 |
| 18 | *Marinobacter daqiaonensis* | 99.5% | 68.87 |
| 19 | *Ketobacter alkanivorans* | 99.5% | 84.04 |
| 20 | *Pseudomonas zhaodongensis* | 99.1% | 76.80 |
| 21 | *Ketobacter sp.* | 99.6% | 77.35 |
| 22 | *Burkholderia sp.* | 99.1% | 76.69 |
| 23 | *Halopseudomonas xiamenensis* | 98.6% | 71.37 |
| 24 | *Halopseudomonas pertucinogena* | 99.5% | 80.15 |
| 25 | *Piscinibacter sp.* | 100% | 85.09 |
| 26 | *Noviherbaspirillum aridicola* | 99.6% | 77.48 |
| 27 | *Pseudomonas nanhaiensis* | 99.1% | 65.87 |

**Supplementary Table 4: $T_M$ Values:** The Tm values for each protein were generated using GraSR, which facilitates rapid and precise comparison of protein structures via contrastive graph neural networks. This study offers information into the structural stability and similarity of the proteins.

| No | Bacteria name | $T_M$ Value |
|---|---|---|
| 1 | *Ideonella sakaiensis* | 0.93088 |
| 2 | *Acidovorax delafieldii* | 0.93634 |
| 3 | *Candidatus Pseudomonas excrementavium* | 0.93765 |
| 4 | *Ectothiorhodospiraceae bacterium WFHF3C12* | 0.93202 |
| 5 | *Massilia sp.* | 0.91205 |
| 6 | *Cytophagales bacterium* | 0.93569 |
| 7 | *Aquabacterium sp.* | 0.93524 |
| 8 | *Pseudomonas sp.* | 0.9313 |
| 9 | *Oleispira antarctica* | 0.93409 |
| 10 | *Proteobacteria bacterium* | 0.92082 |
| 11 | *Rhizobacter sp.* | 0.93569 |
| 12 | *Candidatus Muproteobacteria bacterium* | 0.92477 |
| 13 | *Gammaproteobacteria bacterium HGW-Gammaproteobacteria-6* | 0.93078 |
| 14 | *Halopseudomonas bauzanensis* | 0.93182 |
| 15 | *Schlegelella brevitalea* | 0.92467 |
| 16 | *Rhizobacter gummiphilus* | 0.92479 |
| 17 | *Halopseudomonas litoralis* | 0.93182 |
| 18 | *Halopseudomonas formosensis* | 0.93343 |
| 19 | *Marinobacter daqiaonensis* | 0.93394 |
| 20 | *Ketobacter alkanivorans* | 0.92948 |
| 21 | *Pseudomonas zhaodongensis* | 0.91174 |
| 22 | *Ketobacter sp.* | 0.90473 |
| 23 | *Burkholderia sp.* | 0.9179 |
| 24 | *Halopseudomonas xiamenensis* | 0.93133 |
| 25 | *Halopseudomonas pertucinogena* | 0.93394 |
| 26 | *Piscinibacter sp.* | 0.93569 |
| 27 | *Noviherbaspirillum aridicola* | 0.92944 |
| 28 | *Pseudomonas nanhaiensis* | 0.93132 |

**Supplementary Table 5:** Position and sequences of the connecting loops showing higher RMSF values.

| **RgPETase** | | |
|---|---|---|
| Position | 200-210 | 230-246 |
| Sequence | NDSIAPNSSHS | GSHSCANSGNSDAGLIG |

| **HpPETase** | | | | | | | | |
|---|---|---|---|---|---|---|---|---|
| Position | 68-76 | 82-90 | 98-108 | 123-129 | 145-165 | 214-221 | 244-250 | 284-292 |
| Sequence | SSLVSGFGGG | PYGVEGTMG | FVSAESSIDWW | NTGFDQP | VEQNTSRTSPVQGMIDINRL | SDLIAPVR | NGSHYCA | LCGPSHTSD |

| **HbPETase** | | | | | | |
|---|---|---|---|---|---|---|
| Position | 67-77 | 83-88 | 98-108 | 123-129 | 140-164 | 284-293 |
| Sequence | SGLVSGFGGG | GTTGTM | FVSAESSIEWW | NTGFDQP | ALDYLVSQNTSRTSPVNGMIDTER | GPNHESDRNI |

| **KsPETase** | | | | | | | |
|---|---|---|---|---|---|---|---|
| Position | 63-73 | 90-98 | 101-110 | 148-155 | 163-172 | 217-227 | 250-264 |
| Sequence | GPYSVRTQNVS | TNAGTNMGA | VPGFVSYENS | ALDLLVSE | SGLVDPSRL | IIACEADVVAP | GGHSFCANSGYSDED |

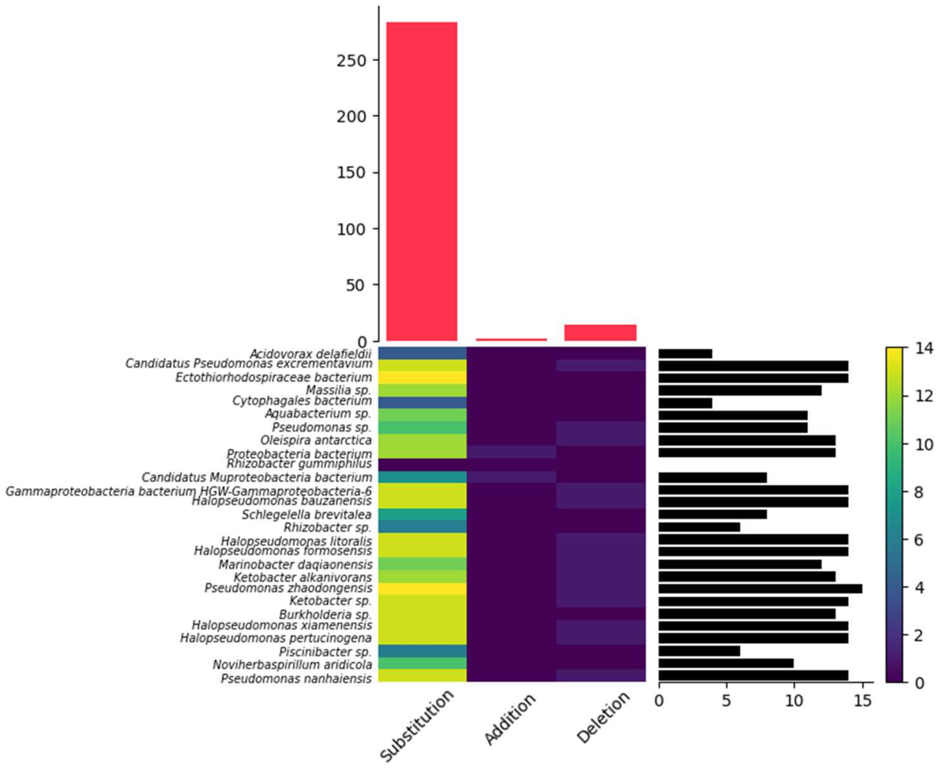


**Supplementary figure S1** - Diversity of AXE domain on the basis of substitution, addition and deletion events.

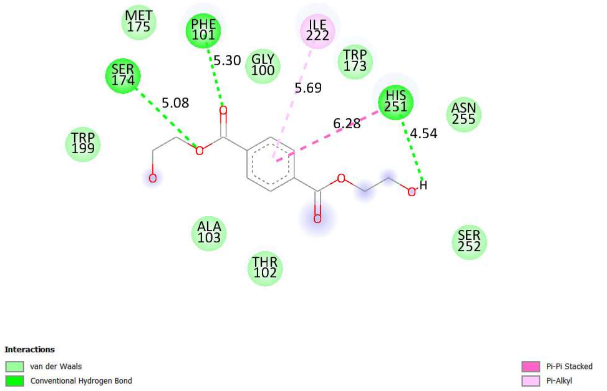


**(a)** – *Acidovorax delafieldii*

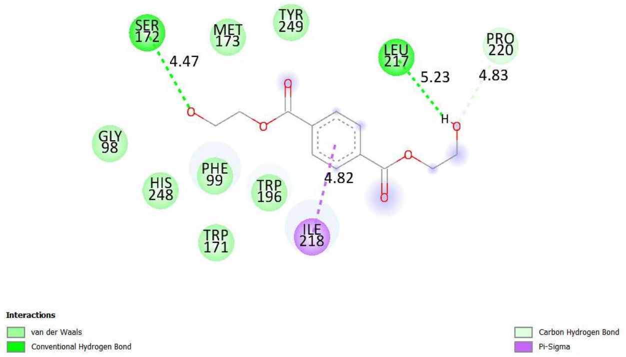


**(b)** – *Candidatus Pseudomonas excrementavium*

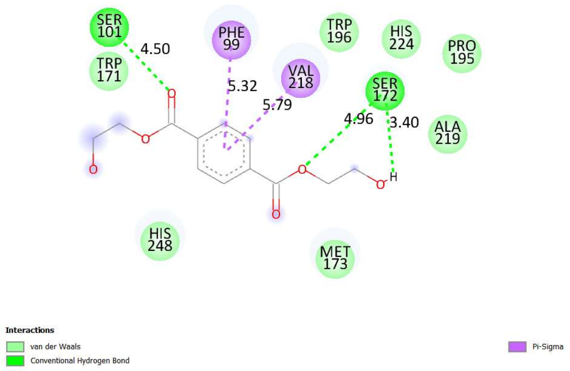


**(c)** - *Pseudomonas nanhaiensis*

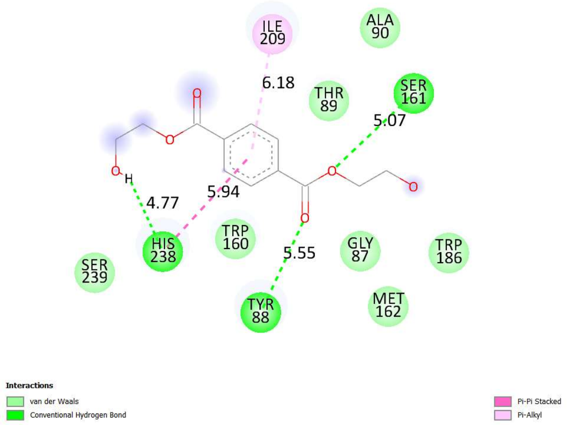


**(d)** - *Piscinibacter sp.*

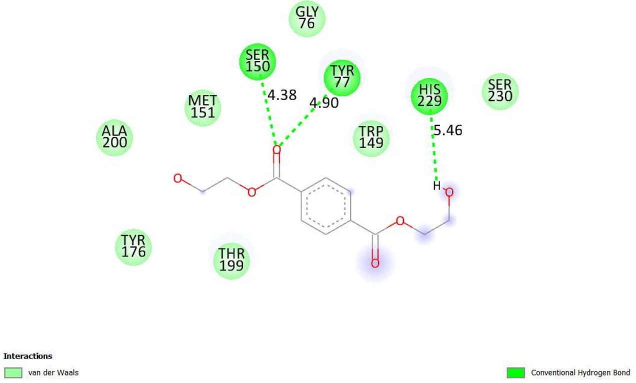


**(e)** - *Candidatus Muproteobacteria* bacterium

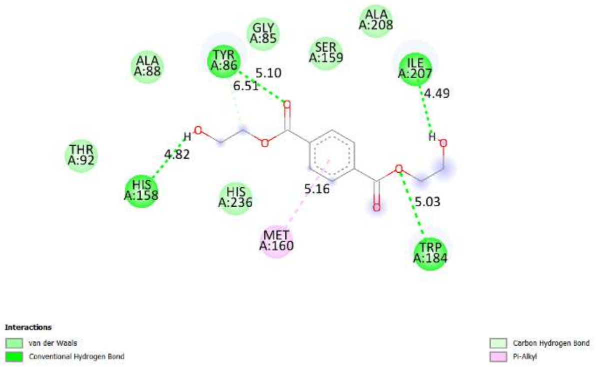


**(f)** – *Aquabacterium sp.*

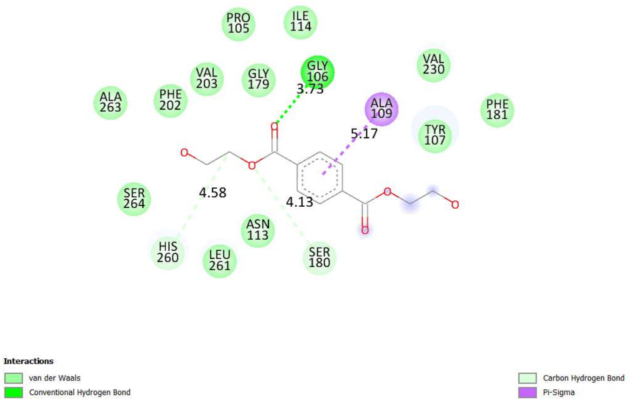


**(g) -** *Burkholderia sp.*

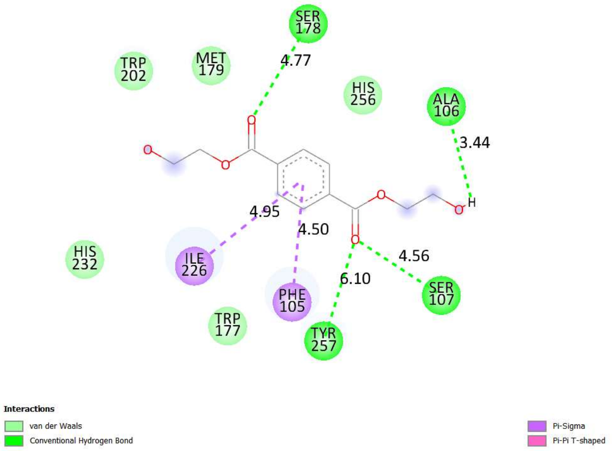


**(h)** - *Marinobacter daqiaonensis*

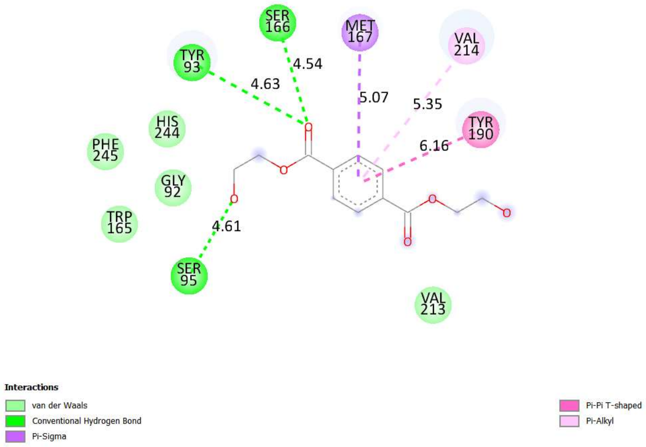


**(i)** - *Pseudomonas zhaodongensis*

**Supplementary figure S2:** The protein-ligand interaction picture depicts the binding interactions between the ligand and particular amino acid residues in the protein's active region. **a)** *Acidovorax delafieldii* **(b)** *Candidatus Pseudomonas excrementavium* **(c)** *Pseudomonas nanhaiensis* **(d)** *Piscinibacter sp.* **(e)** *Candidatus Muproteobacteria* bacterium **(f)** *Aquabacterium sp.* **(g)** *Burkholderia sp.* **(h)** *Marinobacter daqiaonensis* **(i)** *Pseudomonas zhaodongensis*

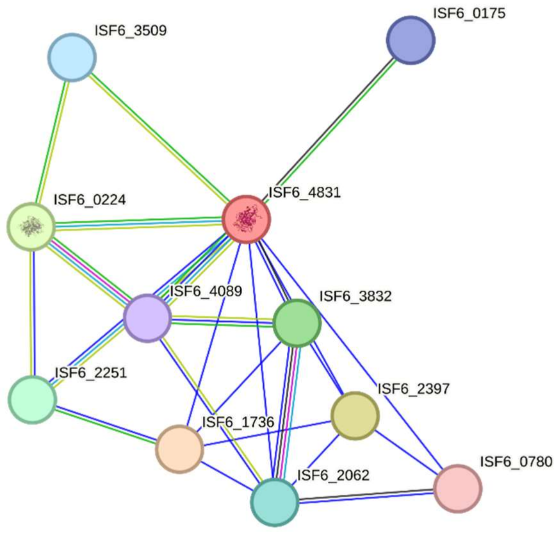


**Supplementary figure S3:** The strong interaction between PETase (ISF6_4831) and MHETase (chlorogenate esterase - ISF6_0224) highlights their collaborative function in PET biodegradation. This association is additionally reinforced by a complex network of interactions with other proteins, emphasising the metabolic pathways and supplementary enzymes that may be implicated in PET breakdown and assimilation. The STRING analysis also finds supplementary proteins (Hypothetical proteins) that may have regulatory or supportive functions in this pathway, suggesting a complex and well-coordinated system for PET degradation inside the organism.

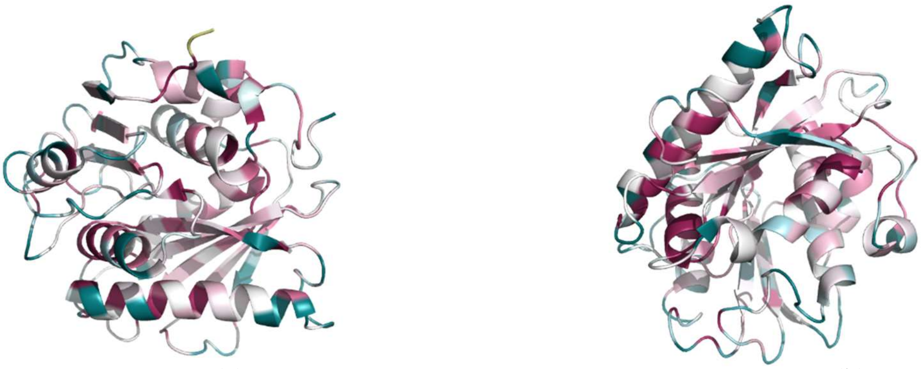

(a) (b)

**Supplementary figure S4** – Conserved regions of IsPETase (a) Anterior face, (b) Posterior face.

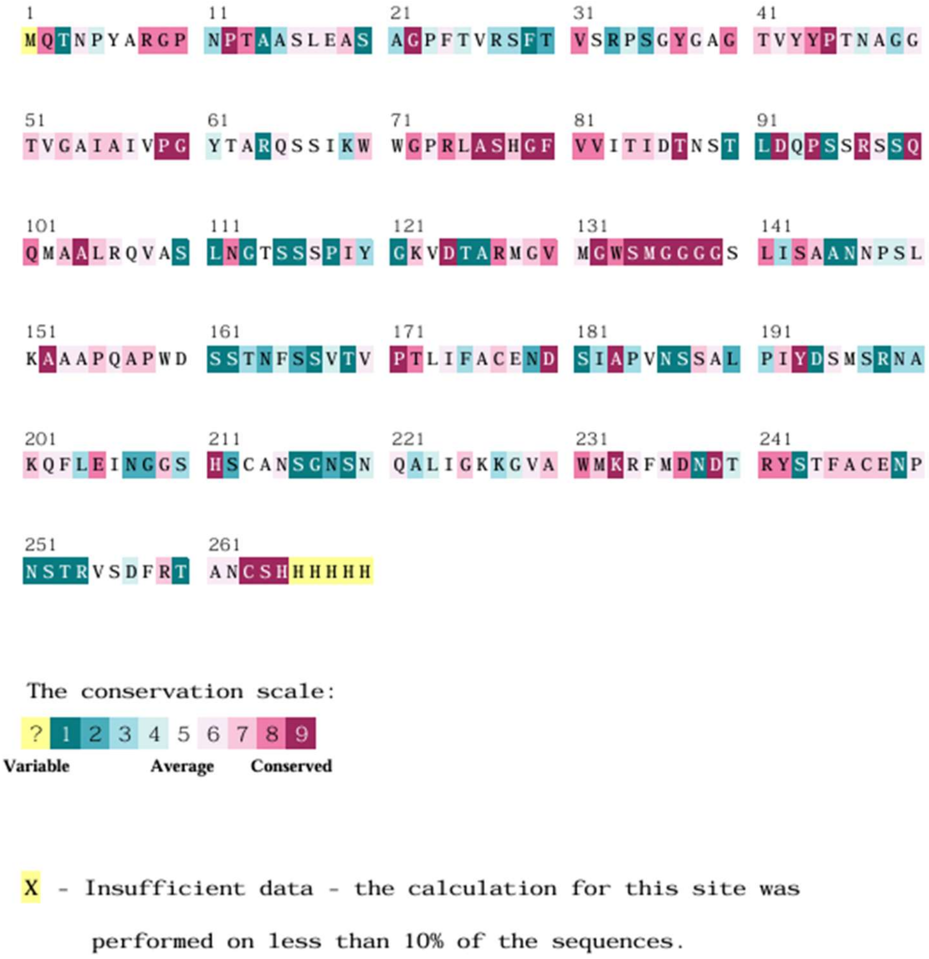


**Supplementary figure S5** – The conserved regions of PETase were analysed using the ConSurf-db tool. Highly conserved sections are indicated by a pink to red colour gradient, while variable regions are represented by a light green to dark green colour gradient.

```
WP_054022242.1  ............................................................
MBX3625601.1    ............................................................
MBC7956640.1    ............................................................
BAB86909.1      ............................................................
WP_047194864.1  ............................................................
WP_085749238.1  ............................................................
MBI3384080.1    ............................................................
MBC7701646.1    ............................................................
OGI66810.1      ............................................................
MBU1194958.1    .................MKFIIALMMSICLMFAAVPAWSAQQTILQESFNSSLGSFTSAG
WP_220809760.1  MKNKNPTLPLLSASPLPGAALAKTGLAVAAMLCSFAA.QAQVAIFQESFSSGLGGFTATG
WP_175819463.1  ............................................................
WP_156871850.1  ............................................................
WP_101893885.1  ............................................................
MBQ0729274.1    ............................................................
WP_122164848.1  ............................................................
MBL4611384.1    ............................................................
WP_188635463.1  ............................................................
WP_090538641.1  ............................................................
WP_036989706.1  ............................................................
HIZ51049.1      ............................................................
WP_090272969.1  ............................................................
WP_185266504.1  ............................................................
PKM05449.1      ............................................................
WP_223655417.1  ............................................................
WP_092009807.1  ............................................................
MBA1146895.1    ............................................................
consensus>50    ............................................................

WP_054022242.1  ............................................................
MBX3625601.1    ............................................................
MBC7956640.1    ............................................................
BAB86909.1      ............................................................
WP_047194864.1  ............................................................
WP_085749238.1  ............................................................
MBI3384080.1    ............................................................
MBC7701646.1    ............................................................
OGI66810.1      ............................................................
MBU1194958.1    SVSVGSSGALMRGSAYSTDGRITSAAVSTQGLTNIVLSFTSATSGLDLGEAAIVSYSTNG
WP_220809760.1  TVGTSTGAARMEGCYGCTDGAITSGAISTQGFTGLSLSFDRVTSGLDSGEAGIAEYSTNG
WP_175819463.1  ............................................................
WP_156871850.1  ............................................................
WP_101893885.1  ............................................................
MBQ0729274.1    ............................................................
WP_122164848.1  ............................................................
MBL4611384.1    ............................................................
WP_188635463.1  ............................................................
WP_090538641.1  ............................................................
WP_036989706.1  ............................................................
HIZ51049.1      ............................................................
WP_090272969.1  ............................................................
WP_185266504.1  ............................................................
PKM05449.1      ............................................................
WP_223655417.1  ............................................................
WP_092009807.1  ............................................................
MBA1146895.1    ............................................................
consensus>50    ............................................................

                                 1       10        20        30
WP_054022242.1  .................MNFPRASRLMQAAVLGGLMAVSAAATAQTNP............
MBX3625601.1    .................MNFPRASRLMQAAVLGGLMAVSAAATAQTNP............
MBC7956640.1    ................MTTTHGIARFAKASLVIGALMLSAAASAQ.NP............
BAB86909.1      ...MHLPRSRWDIPFKEETTMTHHFSVRALLAAGALLASAAVSAQTNP............
WP_047194864.1  ...............MPPDCVLPRRLAAAALLASATLVPLSAAAQTNP............
WP_085749238.1  ....................MKLNRLFQVACLAATLVTATAASA................
MBI3384080.1    ..................MPLQLTNRYGLAALACAALMTLGGHAQAQS............
MBC7701646.1    ..............MTHTTHSKFSRLLQVSLGVSALALSLAAGAQ..S............
OGI66810.1      ..............................MSLQGTGGTTEPPPTGNP............
MBU1194958.1    SSYTTVSSERYAT.GSFTFNLGTAAANQSQLFLRFAIDASSTYETYAVDNIILTGDGEGG
WP_220809760.1  STWTAVESTRVTTNGRVTFSLPDTAAGQAALRLRFRIDASLSSETYTVDNVQLTGIPGDT
WP_175819463.1  ...............MITWNKLQRMVAHSSGNMRGNVFRKDIVLLVLVFSLPLAGRAQSA
WP_156871850.1  ..............MKKKLLATTTTLLASAMLSASVFALTDPPVDPVDPV.DPTDPPSSD
WP_101893885.1  ..............MKLDILKPLIVTLVAGMLSSAAFAITPSPTPTPDPTPNPSPDPGSC
MBQ0729274.1    ..............MNKSILKKLSFGTSVLLVSMNALSWTPSPTPNPDPIPDPTPCQDDC
WP_122164848.1  ..............MKLKAYLARITSLVTVSLLASSLAYA.........APGPSAPCADC
MBL4611384.1    .........................MFSASTFAYNPAPPPPSPTPTPTPGPLPG..Q..P
WP_188635463.1  .............MINTFLPKSLASVLAAGALLLSSAAMANNPPVDP.PPG........G
WP_090538641.1  .............MINNIFPKSLLSMIAAGALLMSASAFATNPPADP.PPG........G
WP_036989706.1  .............MINKNLSQSLLAMMAAGALLLSSSAFAVNPPTDG.PTD........P
HIZ51049.1      .............MINKNLSQSLLAMMAAGALLLSSSAFATNPPTDGSPTD........P
WP_090272969.1  .............MINKNLPSSLLSMLAAGALMLSTSAFATNPPVDD.PSD........P
WP_185266504.1  .............MINKTLSNSLLSMLAAGALLLSTSVMATNPPVDE.PVD........P
PKM05449.1      .............MKNHSFAKTVLSALAAGALLFSAATMASNPTPTPTPTPTPTPTP..G
WP_223655417.1  .............MNNKTFHKSALSLLAASALLFSASAMSSNPTPPP.PTD.....P..G
WP_092009807.1  .............MFRILGKKTALSLLAAGSLMVSASAFAISGGGDDGGGDNGGGCE..T
MBA1146895.1    .............MDPSLKRALGLVVALTAAMLFSASASAQNPPPDDDPEPCQTDCDDGG
consensus>50    .........................ll.a..ll......a..p.................
```

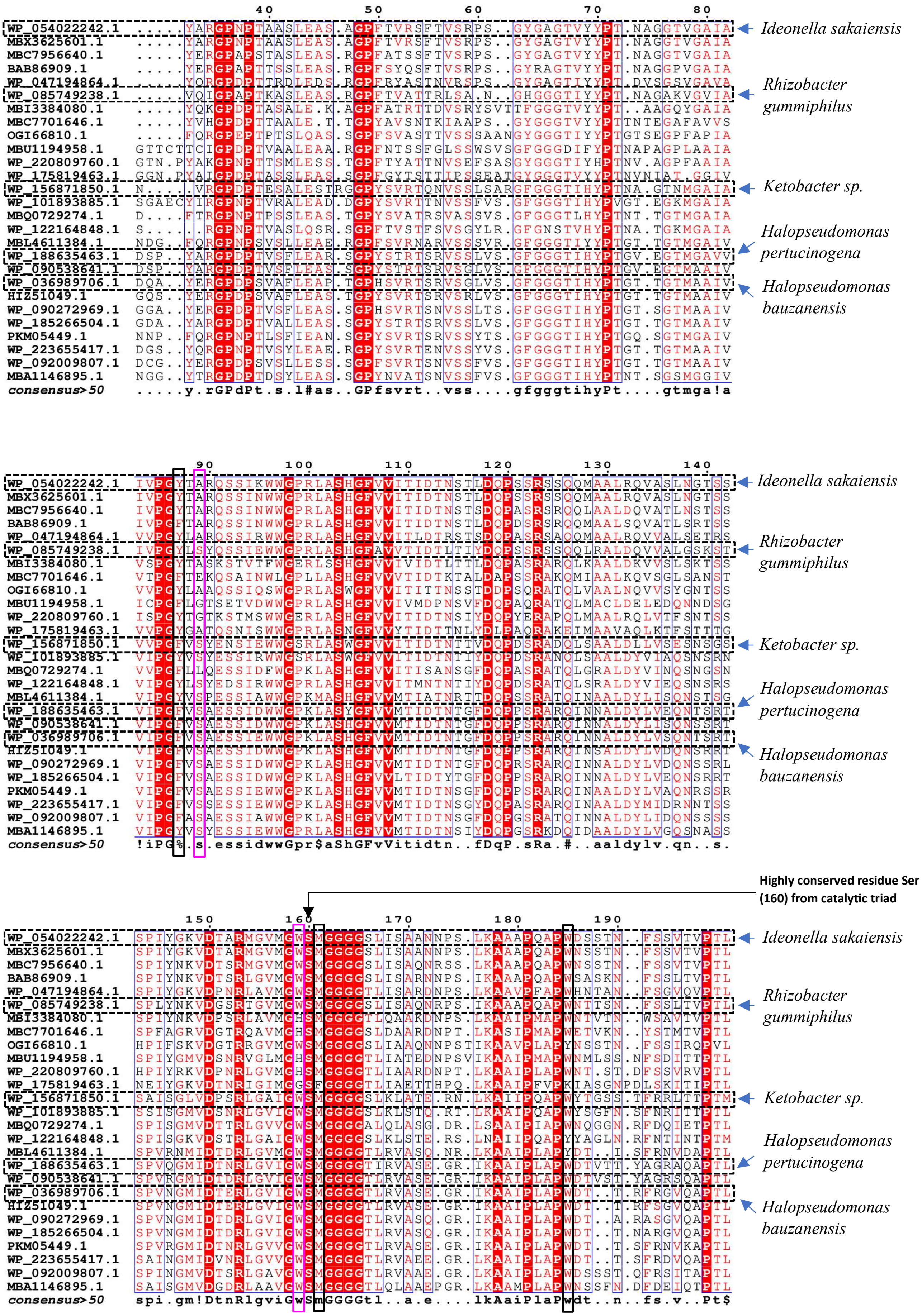
WP_054022242.1
MBX3625601.1
MBC7956640.1
BAB86909.1
WP_047194864.1
WP_085749238.1
MBI3384080.1
MBC7701646.1
OGI66810.1
MBU1194958.1
WP_220809760.1
WP_175819463.1
WP_156871850.1
WP_101893885.1
MBQ0729274.1
WP_122164848.1
MBL4611384.1
WP_188635463.1
WP_090538641.1
WP_036989706.1
HIZ51049.1
WP_090272969.1
WP_185266504.1
PKM05449.1
WP_223655417.1
WP_092009807.1
MBA1146895.1
consensus>50
Ideonella sakaiensis
Rhizobacter gummiphilus
Ketobacter sp.
Halopseudomonas pertucinogena
Halopseudomonas bauzanensis
Highly conserved residue Ser (160) from catalytic triad

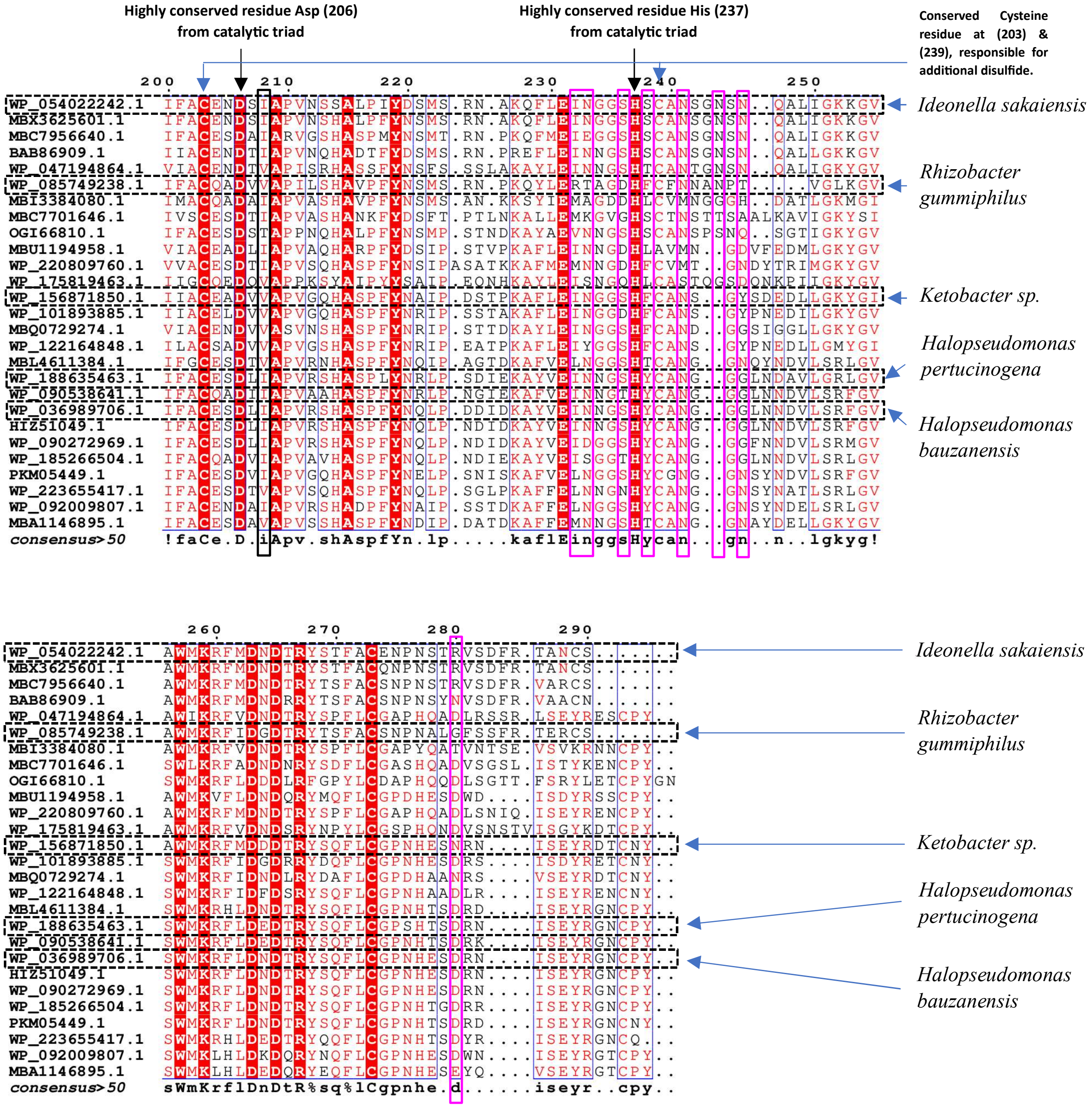


**Supplementary figure S6:** Multiple sequence alignment of 27 prospective PETases executed with CLUSTALW**.** Accession numbers are present on the left side of the image and scientific name details for each accession number is mentioned in supplementary Table 1. Residues of subsite 1 are marked with a black box, while residues of subsite 2 are outlined with a pink box.

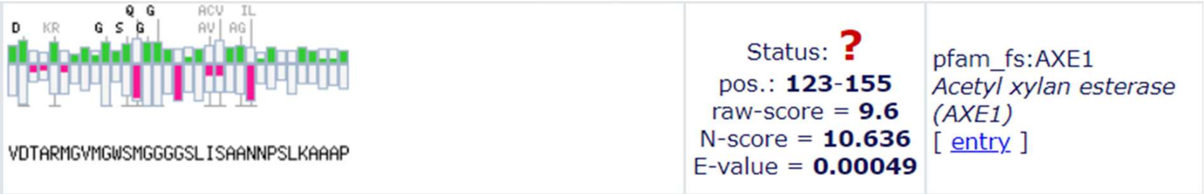


**Supplementary figure S7:** Identified Acetyl xylan esterase (AXE1) family from the Pfam database corresponds to locations 123–155 of the IsPETase sequence, demonstrating a raw score of 9.6 and a significant E-value of 0.00049, indicating a strong alignment with the AXE1 domain.

**AXE domain alignment.**

Serine hydrolase motif (G-X1-S-X2-G)

```
T-COFFEE, Version_11.00 (Version_11.00)
Cedric Notredame
CPU TIME:0 sec.
SCORE=98

BAD AVG GOOD

Chain              VDTARMGVMGWSMGGGGSLISAANNPS-LKAAAP   Chain           :  98
MBX3625601.1       VDTARMGVMGWSMGGGGSLISAANNPS-LKAAAP   MBX3625601.1    :  99
MBC7956640.1       VDTSRMGVMGWSMGGGGSLISAQNRSS-LKAAAP   MBC7956640.1    :  99
WP_201805836.1     VDASRLGVMGWSMGGGGSLISARDNPS-IKAAAP   WP_201805836.1  :  99
MBU1194958.1       VDSNRVGLMGHSMGGGGTLIATEDNPSVIKAAIP   MBU1194958.1    :  97
WP_156871850.1     VDPSRLGAIGWSMGGGGSLKLATE-R-NLKAIIP   WP_156871850.1  :  99
MBC7701646.1       VDGTRQAVMGHSMGGGGSLDAARDNPT-LKASIP   MBC7701646.1    :  99
WP_175819463.1     VDTNRIGIMGGSFGGGGTLIAETTHPQ-LKAAIP   WP_175819463.1  :  92
MBL4611384.1       IDTDRLGVMGWSMGGGGTLRVASQ-N-RLKAAIP   MBL4611384.1    :  98
WP_188635463.1     IDTNRLGVIGWSMGGGGTIRVASE-G-RIKAAIP   WP_188635463.1  :  99
WP_101893885.1     VDSNRLGVIGWSMGGGGSLKLSTQ-R-TLKAAIP   WP_101893885.1  :  99
WP_122164848.1     VDSTRLGAIGWSMGGGGSLKLSTE-R-SLNAIIP   WP_122164848.1  :  99
WP_036989706.1     IDTERLGVIGWSMGGGGTLRVASE-G-RIKAAIP   WP_036989706.1  :  99
MBQ0729274.1       VDTTRLGVVGWSMGGGGALQLASG-DR-LSAAIP   MBQ0729274.1    :  98
BAB86909.1         VDTSRLGVMGWSMGGGGSLISARNNPS-IKAAAP   BAB86909.1      :  99
WP_047194864.1     VDPNRLAVMGWSMGGGGTLISARDNPS-LKAAVP   WP_047194864.1  :  99
WP_085749238.1     VDGSRTGVMGWSMGGGGSLISAQNRPS-IKAAAP   WP_085749238.1  :  99
OGI66810.1         VDGTRRGVMGWSMGGGGSLIAAQNNPSTIKAAVP   OGI66810.1      :  97
WP_220809760.1     VDPNRLGVMGHSMGGGGTLIAARDNPT-LKAAIP   WP_220809760.1  :  99
MBI3384080.1       VDPSRLAVMGHSMGGGGTLQAAKDNPS-LKAAIP   MBI3384080.1    :  99
WP_092009807.1     VDTSRLGAVGWSMGGGGTLRVAAE-G-RLKAAIP   WP_092009807.1  :  99
PKM05449.1         IDTNRLGVVGWSMGGGGTLRVASE-G-RIKAAIP   PKM05449.1      :  99
HIZ51049.1         IDTERLGVIGWSMGGGGTLRVASE-G-RIKAAIP   HIZ51049.1      :  99
MBA1146895.1       VDGDRLAAVGWSMGGGGTLQLAAE-PGRLKAAMP   MBA1146895.1    :  97
WP_090272969.1     IDTDRLGVIGWSMGGGGTLRVASQ-G-RIKAAIP   WP_090272969.1  :  98
WP_090538641.1     IDTDRLGVIGWSMGGGGTLRVASE-G-RIKAAIP   WP_090538641.1  :  99
WP_223655417.1     IDVNRLGVVGWSMGGGGTLRVAEE-G-RLKAAIP   WP_223655417.1  :  99
WP_185266504.1     IDTDRLGVIGWSMGGGGTLRVASQ-G-RIKAAIP   WP_185266504.1  :  98

cons               :*  * . :* *:****::         :.*  *   cons            :   9
```

**Supplementary figure S8:** Alignment of AXE domain: The alignment shows a score of 98, signifying excellent conservation among the sequences. The accession numbers of the proteins are displayed on the left, and conserved residues are indicated below the alignment with symbols (* , :, and .), denoting entirely conserved, highly similar, and moderately similar areas, respectively. The majority of the alignment is marked in red/pink indicating highly conserved regions.

**Ligand RMSD**

The ligand RMSD indicates whether the ligand maintains its original binding conformation or undergoes substantial conformational alterations. Comparatively low and consistent RMSD values (ranging from 0.1 to 0.2 nm) with minimal variations indicate stable binding conformations within the protein binding pocket. Minor fluctuations appear due to slight conformational adjustments of the ligand.

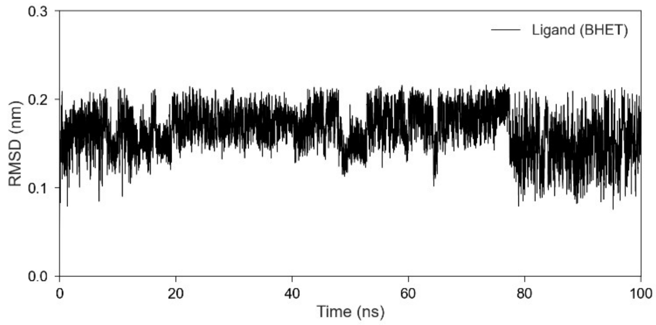


**(a)**

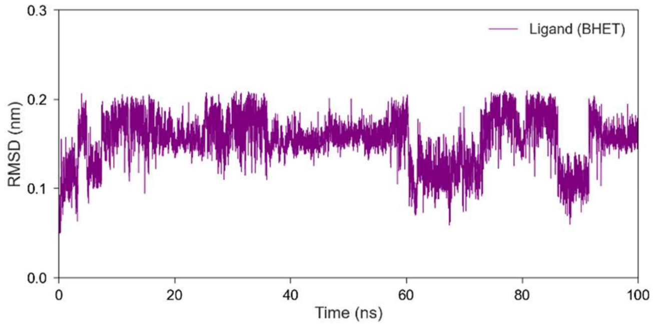


**(b)**

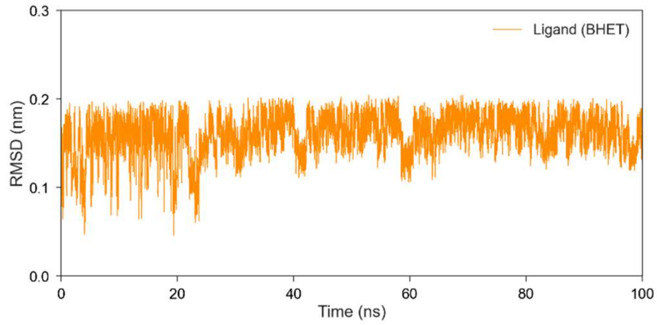


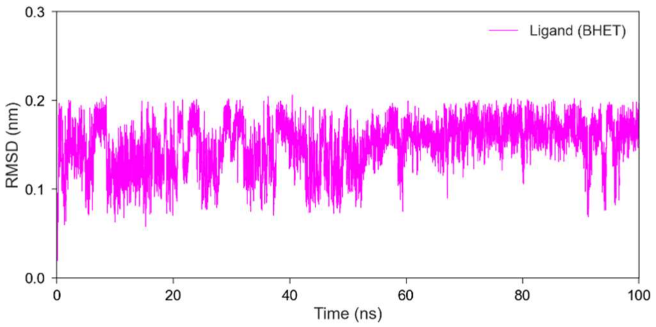


**(c)** **(d)**

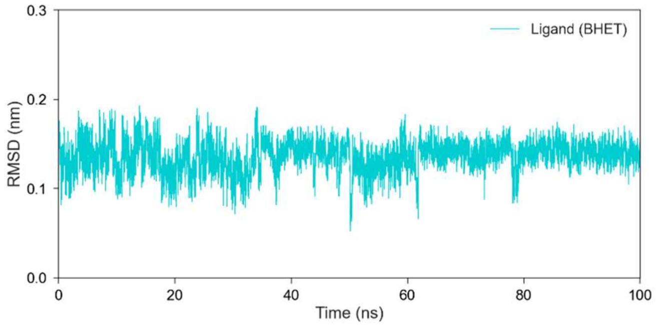


**(e)**

**Supplementary figure S9:** Ligand RMSD indicates the conformational stability of the ligand. The ligand binding across all five proteins exhibits a ligand RMSD ranging from 0.1 to 0.2 nm, indicating a low value that reflects a very stable binding configuration throughout the simulation. a) PETase from *Ideonella sakaiensis* (black colour) b) PETase from *Rhizobacter gummiphilus* (purple) c) PETase from *Halopseudomonas pertucinogena* (orange) d) PETase from *Halopseudomonas bauzanensis* (pink) e) PETase from *Ketobacter sp.* (cyan)
